# Immunoproteasome Deficiency Impairs Microglial Clearance and Worsens Tau and Amyloid Pathology

**DOI:** 10.64898/2026.07.14.738427

**Authors:** Malavika Srikanth, Shan Jiang, Steven M. Wellman, Shrita Sarkar, Daniella E. Lorman, Timothy Lantin, Avery M. Runyan, Mayank Kumar, Eric Sydney, Helen Y. Figueroa, Mu Yang, Qi Wang, Natura Myeku

## Abstract

Immunoproteasome induction is prominent in Alzheimer’s disease (AD), but whether it protects proteostasis or amplifies neuroinflammation remains unresolved. Here, we generated immunoproteasome-deficient PS19 tauopathy and APP/human tau double-knock-in mice by crossing each disease model with L7M1 mice lacking two immunoproteasome catalytic subunits. Immunoproteasome deficiency increased phospho-tau burden, exacerbated amyloid-β pathology and heightened microglial reactivity without suppressing constitutive 26S proteasome activity. In primary microglia and longitudinal two-photon imaging, immunoproteasome-deficient microglia engaged and engulfed tau aggregate–bearing material but failed to resolve internalized cargo, revealing a post-engulfment degradative checkpoint. Single-nucleus transcriptomics identified a remodeled P2ry12^low^/Trem2^high^ microglial state with impaired phagolysosomal and mitochondrial programs. Reanalysis of human single-nucleus transcriptomic datasets showed that reduced microglial immunoproteasome expression was associated with cargo-processing gene-program changes similar to those observed in immunoproteasome-deficient mouse microglia. Together, these findings identify immunoproteasome biogenesis as a protective glial stress response that supports microglial aggregate clearance in AD.

---

The immunoproteasome is an inducible proteasome variant that expands cellular proteolytic capacity during inflammatory and oxidative stress. It is formed when the canonical β1, β2, and β5 catalytic subunits of the 20S core proteasome are replaced by the inducible β1i/LMP2, β2i/MECL-1, and β5i/LMP7 subunits in response to cytokines such as interferon-γ and tumor necrosis factor-α, as well as stress-responsive inflammatory pathways ^1–4^. Although originally defined by its role in MHC class I antigen presentation, the immunoproteasome also exhibits altered cleavage-site specificity and enhanced capacity to remove oxidatively damaged or structurally abnormal proteins, thereby linking immune activation to proteostasis ^5–8^. This dual role is particularly relevant in the central nervous system (CNS), where glial cells must coordinate inflammatory responses with the clearance of damaged proteins, aggregate-prone substrates and neuronal debris during aging and neurodegenerative disease.

In AD, proteasome remodeling has been observed in both human tissue and mouse models. Reactive glia surrounding amyloid plaques show increased immunoproteasome expression and activity ^9–11^, while recent multi-omics work has shown early impairment of constitutive proteasome programs and the stage-dependent emergence of immunoproteasome components across Braak stages, particularly in glial cells ^12^. These observations support the idea that immunoproteasome induction may represent a compensatory response to declining neuronal proteostasis and increasing proteotoxic burden in the inflammatory AD brain. However, whether this response is protective or pathogenic remains unresolved. Immunoproteasome activity can intersect with inflammatory signaling, and pharmacological inhibition or genetic disruption of immunoproteasome components has been reported to attenuate cytokine responses or improve selected phenotypes in some AD-relevant contexts^9–11,13^. Conversely, studies of oxidative-stress proteostasis indicate that immunoproteasomes can promote the clearance of damaged or misfolded proteins and protect cells from proteotoxic injury ^5–7^. Thus, whether immunoproteasome induction in AD primarily amplifies neuroinflammation, preserves glial degradative capacity, or reflects a context-dependent balance between these functions remains unknown.

This question is especially important for microglial handling of pathological tau and amyloid-associated material. When intracellular processing is efficient, this response may limit persistence of pathogenic protein species and restrict their intercellular spread. Under chronic inflammatory or oxidative stress, however, microglia may remain engaged with pathological cargo while failing to complete degradation. In this setting, uptake alone may not indicate successful clearance; instead, incompletely processed tau or amyloid-associated material may persist intracellularly, sustain microglial stress responses, or re-enter the extracellular space^14–16^. At the same time, immunoproteasome-dependent antigen-processing capacity through MHC class I pathways may intersect with T-cell-mediated inflammatory circuits in the CNS^17–19^. Defining how inducible immunoproteasome subunits influence microglial cargo processing is therefore essential for understanding whether immunoproteasome induction is adaptive or maladaptive in AD.

To determine how immunoproteasome function shapes AD-related pathology in vivo, we used the L7M1 mouse line, which is deficient in the inducible immunoproteasome catalytic subunits β2i/MECL-1/PSMB10 and β5i/LMP7/PSMB8^13,20^. We crossed L7M1 mice with two complementary AD-relevant models: the PS19 tauopathy line, which expresses human P301S tau and develops progressive tau pathology with synaptic dysfunction and robust glial activation^21^, and an amyloid-tau double knock-in (dKI) model combining the App-NL-G-F mutant allele with a humanized MAPT allele expressed from the endogenous locus^22,23^. Using histopathology, biochemical assays, behavioral phenotyping, in vitro live-cell imaging, in vivo two-photon microscopy, mouse single-nucleus transcriptomics, and reanalysis of human single-nucleus transcriptomic datasets, we show that immunoproteasome deficiency exacerbates tau and amyloid pathology and shifts microglia into an activated but inefficient cargo-processing state. Immunoproteasome-deficient microglia engage and engulf tau-bearing material but fail to efficiently resolve internalized cargo, revealing a post-engulfment degradative checkpoint. Together, these findings identify the immunoproteasome as a protective, stress-adaptive component of glial proteostasis that supports microglial aggregate clearance in AD-relevant brain environments.

## RESULTS

### Immunoproteasome deficiency exacerbates tau and amyloid-β accumulation and glial activation in PS19 and APP/hTau dKI mice

To determine whether immunoproteasomes influence tauopathy and glial responses, we crossed L7M1 mice with PS19 mice, which develop robust tau pathology by approximately 6 months of age ^20,21^. In hippocampal sections immunostained for pathological hyperphosphorylated tau (pS202/pT205; AT8) and the microglial marker Iba1, L7M1/PS19 mice showed increased tau inclusions relative to PS19 controls, as reflected by the percentage area of AT8-positive signal (**Fig. 1A,B; P < 0.0001**). Microglial reactivity was also increased, with greater Iba1-positive area (**Fig. 1A,C; P < 0.0001**) and elevated numbers of Iba1-positive cells (**Fig. 1A,D; P < 0.001**). Low-magnification images are shown in **Supplementary Fig. 1A,B.** Consistent with the histological findings, immunoblotting revealed elevated pS202/pT205 tau in L7M1/PS19 brain lysates compared with PS19 controls (**Fig. 1E**), and densitometric quantification confirmed a significant increase in phospho-tau levels (**Fig. 1F; P < 0.01**). Native in-gel activity assays detected no change in constitutive 26S proteasome activity between groups (**Supplementary Fig. 2A-D**). Co-staining for the microglial/myeloid activation-associated marker CD11b further showed increased CD11b immunoreactivity in L7M1/PS19 brains (**Fig. 1G,H; P < 0.0001**) and a greater number of CD11b-positive cells (**Fig. 1G,I; P < 0.01**). Thus, in a tau-driven disease context, immunoproteasome deficiency increased pathological phospho-tau burden in parallel with enhanced microglial reactivity, without detectable suppression of constitutive 26S proteasome activity.

**Figure 1.**
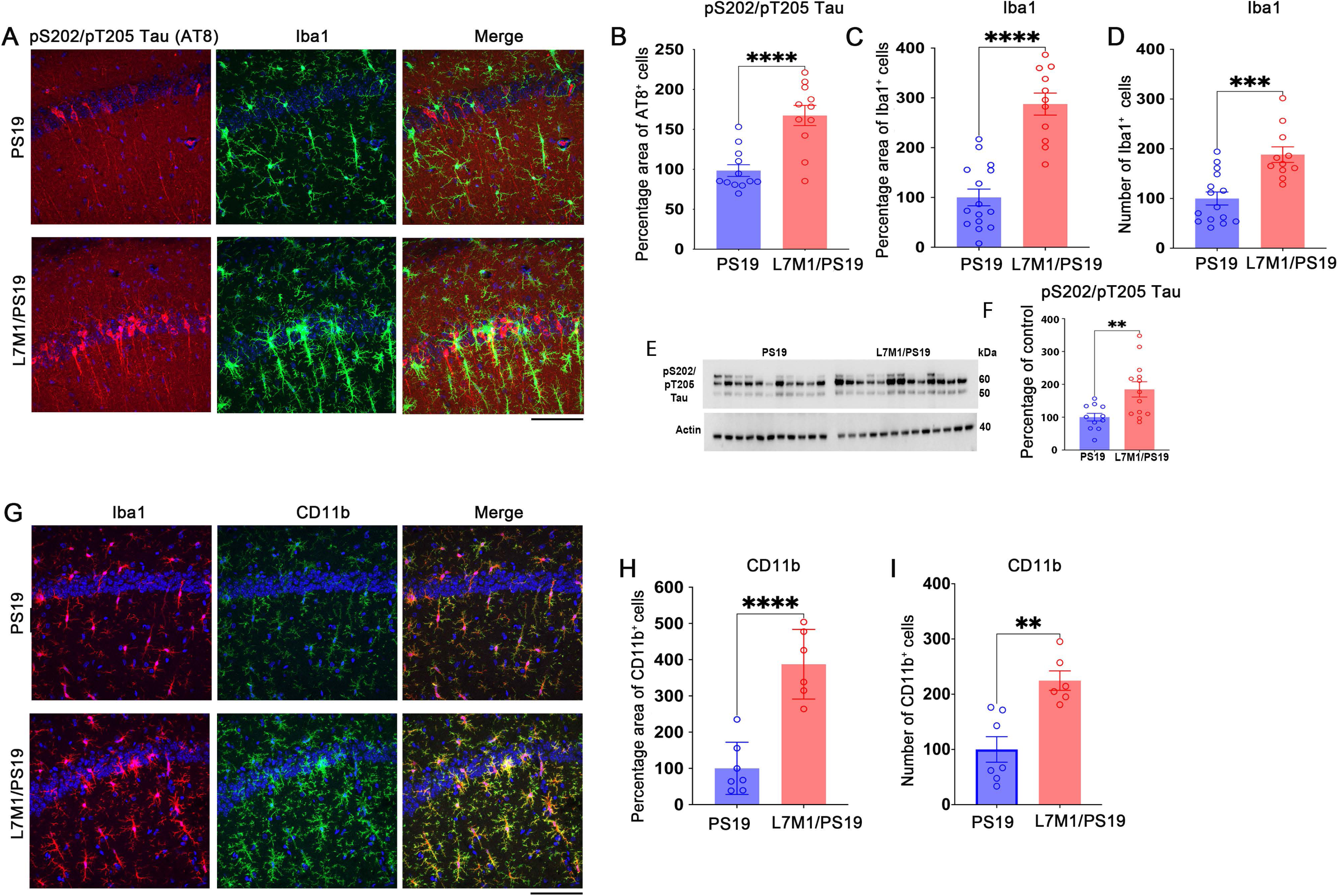
Immunoproteasome deficiency exacerbates phospho-tau pathology and microglial activation in PS19 tauopathy mice. (**A**) Representative immunofluorescence images of hippocampal sections from PS19 and L7M1/PS19 mice stained for phosphorylated tau using pS202/pT205 tau antibody AT8 (red), the microglial marker Iba1(green) and DAPI. Merged images show increased phospho-tau burden and microglial reactivity in L7M1/PS19 mice. (**B**) Quantification of AT8-positive phospho-tau burden, shown as percentage area of AT8-positive staining. (**C**) Quantification of Iba1-positive microglial staining area and (**D**) number of Iba1-positive cells. (**E**) Representative immunoblot of pS202/pT205 tau in PS19 and L7M1/PS19 brain lysates, with actin shown as a loading control. (**F**) Quantification of pS202/pT205 tau immunoblot signal, expressed as percentage of PS19 control. (**G**) Representative immunofluorescence images of hippocampal sections stained for Iba1 (red), CD11b (green) and DAPI. Merged images show increased CD11b-positive microglial activation in L7M1/PS19 mice. (**H**) Quantification of CD11b-positive staining area and (**I**) number of CD11b-positive cells. For quantifications, each plotted circle represents one mouse. For each mouse, four sections were analyzed, with three images acquired per section; the resulting 12 images were averaged to generate one biological replicate per animal. Normality was assessed by Shapiro–Wilk test, and groups were compared using an unpaired two-tailed t-test. Data are shown as mean ± s.e.m. ns, not significant; P values are indicated as **P < 0.01, ***P < 0.001 and ****P < 0.0001. Scale bars, 20 µm.

We next asked whether this relationship extended to an AD-relevant setting in which amyloid and tau-associated pathologies coexist. The APP/hTau dKI model combines human amyloid precursor protein knock-in mutations with humanized MAPT, enabling age-dependent amyloidosis, tau phosphorylation, and chronic microgliosis without transgene overexpression^22,23^. In cortical sections, L7M1/APP/hTau dKI mice showed significant increases in both pS396/pS404 tau (PHF1; **Fig. 2A,B; P < 0.01**) and 6E10-positive amyloid burden (**Fig. 2A,C; P < 0.01**) compared with APP/hTau dKI controls. Iba1-positive area was also increased (**Fig. 2A,D; P < 0.001**), although cortical microglial numbers were not detectably altered (**Fig. 2A,E; n.s**.). A similar pattern was observed in the dentate gyrus, where pS396/pS404 tau (**Fig. 2F,G; P < 0.001**), amyloid burden (**Fig. 2F,H; P < 0.01**), and Iba1-positive area (**Fig. 2F,I; P < 0.001**) were increased in L7M1/APP/hTau dKI mice, whereas microglial numbers were unchanged (**Fig. 2F,J; n.s.**). Low-magnification images are shown in **Supplementary Fig. 1C-E**. As in the PS19 cohort, native in-gel activity assays detected no change in constitutive 26S proteasome activity between groups (**Supplementary Fig. 2E-H**). Immunoblotting corroborated the histological increase in phospho-tau, with densitometry confirming elevated pS396/pS404 tau in L7M1/APP/hTau dKI lysates relative to APP/hTau dKI controls (**Fig. 2K,L; P < 0.05**). These data show that immunoproteasome deficiency exacerbates aggregate pathology and glial activation across both tau-dominant and amyloid-tau disease contexts.

**Figure 2.**
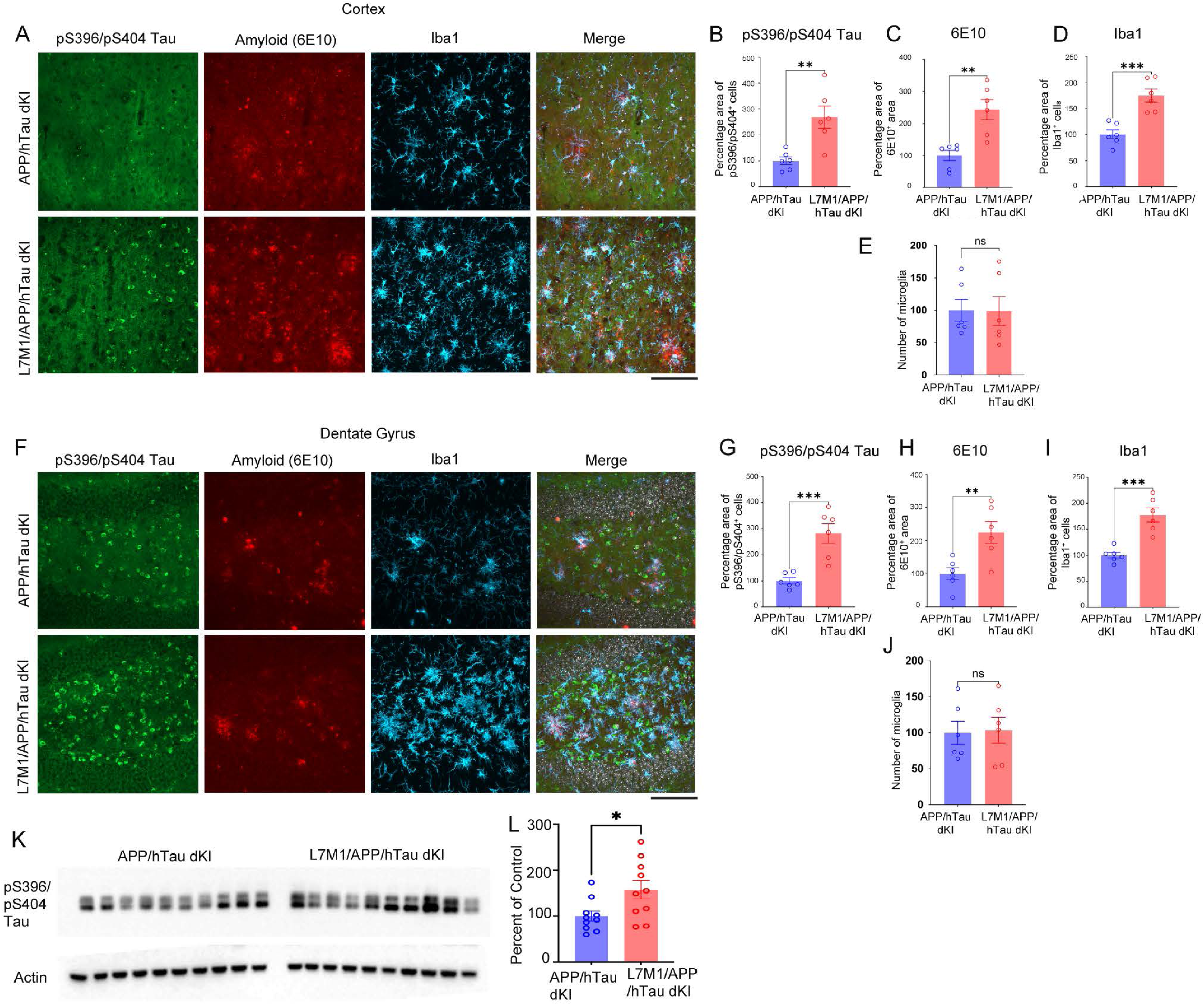
Immunoproteasome deficiency exacerbates tau and amyloid pathology and increases microglial reactivity in APP/hTau dKI mice. (**A**) Representative cortical immunofluorescence images from APP/hTau dKI and L7M1/APP/hTau dKI mice stained for phosphorylated tau at pS396/pS404 (green), amyloid/Aβ using 6E10 (red), and the microglial marker Iba1(cyan). Merged images show increased phospho-tau, amyloid burden, and microglial reactivity in L7M1/APP/hTau dKI mice. (**B)** Quantification in the cortex of the pS396/pS404 tau-positive area (**C**) 6E10-positive amyloid area (**D**) Iba1-positive area (**E**), and microglial number. (**F**) Representative immunofluorescence images from the dentate gyrus of APP/hTau dKI and L7M1/APP/hTau dKI mice stained for pS396/pS404 tau, 6E10 and Iba1. (**G**) Quantification in dentate gyrus of pS396/pS404 tau-positive area. (**H**) 6E10-positive amyloid area. (**I**) Iba1-positive area (**J**) and microglial number. (**K**) Representative immunoblot of pS396/pS404 tau in APP/hTau dKI and L7M1/APP/hTau dKI brain lysates, with actin shown as a loading control. (**L**) Quantification of pS396/pS404 tau immunoblot signal normalized to actin and expressed as percentage of APP/hTau dKI control. For quantifications, each plotted circle represents one mouse. For each mouse, four sections were analyzed, with three images acquired per section; the resulting 12 images were averaged to generate one biological replicate per animal. Normality was assessed by Shapiro–Wilk test, and groups were compared using an unpaired two-tailed t-test Data are shown as mean ± s.e.m. ns, not significant; **\***P < 0.05, **\*\***P < 0.01 and **\*\*\***P < 0.001. Scale bars, 20µm.

### Immunoproteasome deficiency increases swim velocity without altering escape latency

To assess whether immunoproteasome loss influences spatial learning or behavioral state, we subjected PS19 and APP/hTau dKI cohorts, each with or without immunoproteasome deficiency, to Morris water maze testing over 8 consecutive days. Because latency-based measures can be influenced by locomotor drive and stress responsivity, we quantified escape latency alongside swim velocity and distance traveled to distinguish platform acquisition from genotype-dependent differences in swim kinematics^24–26^.

In the PS19 cohort (Non-Tg n = 36; L7M1 n = 38; PS19 n = 25; L7M1/PS19 n = 25), escape latency declined across training in all groups and did not differ significantly among genotypes (**Fig. 3A**). A similar pattern was observed in the APP/hTau dKI cohort (WT n = 14; L7M1 n = 11; APP/hTau dKI n = 13; L7M1/APP/hTau dKI n = 13), where all groups converged on comparable escape latencies over the 8-day training period (**Fig. 3D**). Thus, immunoproteasome deficiency did not measurably alter latency-based acquisition in either disease context.

**Figure 3.**
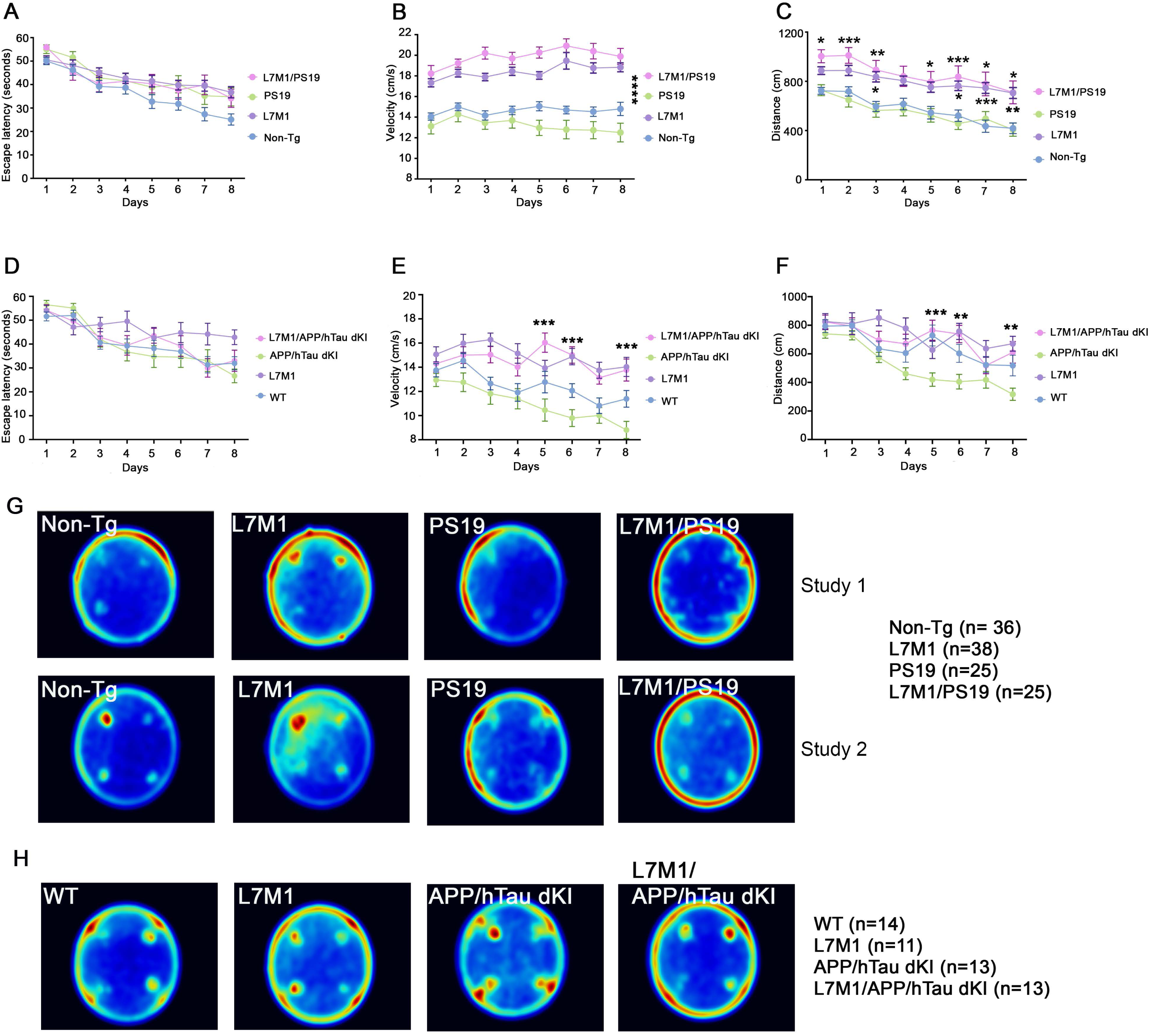
Immunoproteasome deficiency alters Morris water-maze swim behavior without producing a clear latency-based learning deficit. Morris water-maze acquisition in PS19 tauopathy cohorts across 8 days of hidden-platform training. (**A**) Escape latency, (**B**) swim velocity and (**C**) total path length/distance traveled are shown for Non-Tg, L7M1, PS19 and L7M1/PS19 mice. Data are from two combined studies: Non-Tg, n = 36 mice; L7M1, n = 38 mice; PS19, n = 25 mice; L7M1/PS19, n = 25 mice. Morris water-maze acquisition in APP/hTau dKI cohorts across 8 days of hidden-platform training. (**D**) Escape latency, (**E**) swim velocity, and (**F**) total path length/distance traveled are shown for WT, L7M1, APP/hTau dKI and L7M1/APP/hTau dKI mice. WT, n = 14 mice; L7M1, n = 11 mice; APP/hTau dKI, n = 13 mice; L7M1/APP/hTau dKI, n = 13 mice. (**G**) Cumulative occupancy heat maps from PS19 study cohorts, showing search patterns for Non-Tg, L7M1, PS19 and L7M1/PS19 mice across two studies. (**H**) Cumulative occupancy heat maps across 8 days from APP/hTau dKI study cohorts, showing search patterns for WT, L7M1, APP/hTau dKI and L7M1/APP/hTau dKI mice. Warmer colors indicate greater cumulative occupancy. Data are shown as mean ± s.e.m.; each n represents an individual mouse. Behavioral acquisition data across training days were analyzed separately for the PS19 and APP/hTau dKI cohorts using two-way repeated-measures ANOVA, with genotype as the between-subject factor and training day as the repeated-measures factor, followed by Bonferroni-corrected multiple-comparison post hoc tests. Asterisks indicate significant Bonferroni-adjusted post hoc comparisons. ns, not significant; *P < 0.05, **P < 0.01, ***P < 0.001 and ****P < 0.0001.

Swim kinematics diverged more clearly. In the PS19 cohort, L7M1-containing mice showed significantly higher swim velocities than Non-Tg and PS19 controls across training (**Fig. 3B; P < 0.0001**). In the same cohort, distance traveled was separated primarily by immunoproteasome status rather than PS19 genotype: PS19 mice generally tracked Non-Tg controls, whereas L7M1 and L7M1/PS19 mice traveled longer paths at multiple training time points (**Fig. 3C; P values indicated**). In the APP/hTau dKI cohort, L7M1 and L7M1/APP/hTau dKI mice similarly showed increased swim velocity relative to WT and APP/hTau dKI controls from day 5 onward (**Fig. 3E; P < 0.001**), and distance traveled remained elevated in L7M1-containing animals on later training days (**Fig. 3F; P values indicated**). Thus, increased distance was most consistent with the elevated swim velocity/peripheral-search phenotype and did not indicate more efficient platform localization. Cumulative occupancy heat maps further showed redistribution of L7M1-containing genotypes toward peripheral regions of the pool, whereas controls showed more directed occupancy patterns (**Fig. 3G,H**). These findings indicate that immunoproteasome deficiency does not produce a latency-based spatial learning deficit under the conditions tested, but instead causes a hyperkinetic or stress-responsive swim phenotype that can confound interpretation of escape latency as a pure cognitive measure.

### Immunoproteasome-deficient microglia accumulate tau aggregate-bearing cellular material in vitro

Given the increased tau pathology and microglial activation observed in vivo, we next asked whether loss of immunoproteasome function alters the ability of microglia to handle tau aggregate–bearing cellular material. Primary microglia isolated from WT or L7M1 mice were co-cultured with DS9 clone cells, a monoclonal HEK293-derived cell line that stably expresses mutant tau repeat domain-YFP (tauRD-P301L/V337M-YFP) and propagates a defined intracellular tau-aggregate strain used in tau-seeding assays ^27–29^. After 48 h of co-culture, fixed cultures were immunostained for Iba1 to visualize microglia, while DS9-derived tau cargo was detected by YFP fluorescence. Compared with WT microglia, L7M1 microglia showed visibly greater accumulation of DS9-derived tauRD-YFP material (**Fig. 4A**). Quantification confirmed a significant increase in the percentage of DS9 clone cells or DS9-derived material associated with/engulfed by L7M1 microglia relative to WT microglia (**Fig. 4B; P < 0.0001**). These endpoint data indicated that immunoproteasome-deficient microglia accumulate more tau aggregate–bearing cargo, but did not distinguish whether this reflected increased engulfment, impaired intracellular degradation, or both. To distinguish cargo uptake from post-engulfment processing, we performed live-cell imaging in the same DS9–microglia co-culture system. WT and L7M1 microglia were labeled with mCherry and co-cultured with DS9 cells expressing green fluorescent tauRD-YFP cargo. Time-lapse movies were acquired over 22 h, with images collected approximately every 15 min. Representative frames are shown at 30-min intervals over the first 270 min of the recording, while quantification was performed from the full live-imaging movies (**Fig. 4C,D; Supplementary Videos 1 and 2**). In WT microglia, DS9-derived green cargo transiently contacted or entered mCherry-positive microglia and then progressively diminished, consistent with internalization followed by intracellular digestion or clearance (**Fig. 4C and Supplementary Video 1**). In contrast, L7M1 microglia readily contacted and internalized DS9-derived cargo, but the intracellular green signal persisted over time and frequently remained within enlarged intracellular compartments (**Fig. 4D and Supplementary Video 2**). To quantify this phenotype, we calculated digestion efficiency as the ratio of digestion events to engulfment/engagement events. Engulfment/engagement was defined as DS9-derived green cargo contacting, overlapping with, or being internalized by mCherry-positive microglia. Digestion was defined as progressive loss or disappearance of internalized green cargo after microglial uptake. L7M1 microglia showed a significantly reduced digestion-to-engulfment ratio compared with WT microglia (**Fig. 4E; P < 0.01**), despite their ability to engage and internalize DS9-derived material. Thus, immunoproteasome deficiency does not prevent microglial recognition or uptake of tau aggregate–bearing cargo; rather, it impairs efficient post-engulfment intracellular processing. These findings suggest that the increased tau cargo burden observed in L7M1 microglia reflects defective clearance after uptake, resulting in intracellular retention of tau aggregate–bearing material.

**Figure 4.**
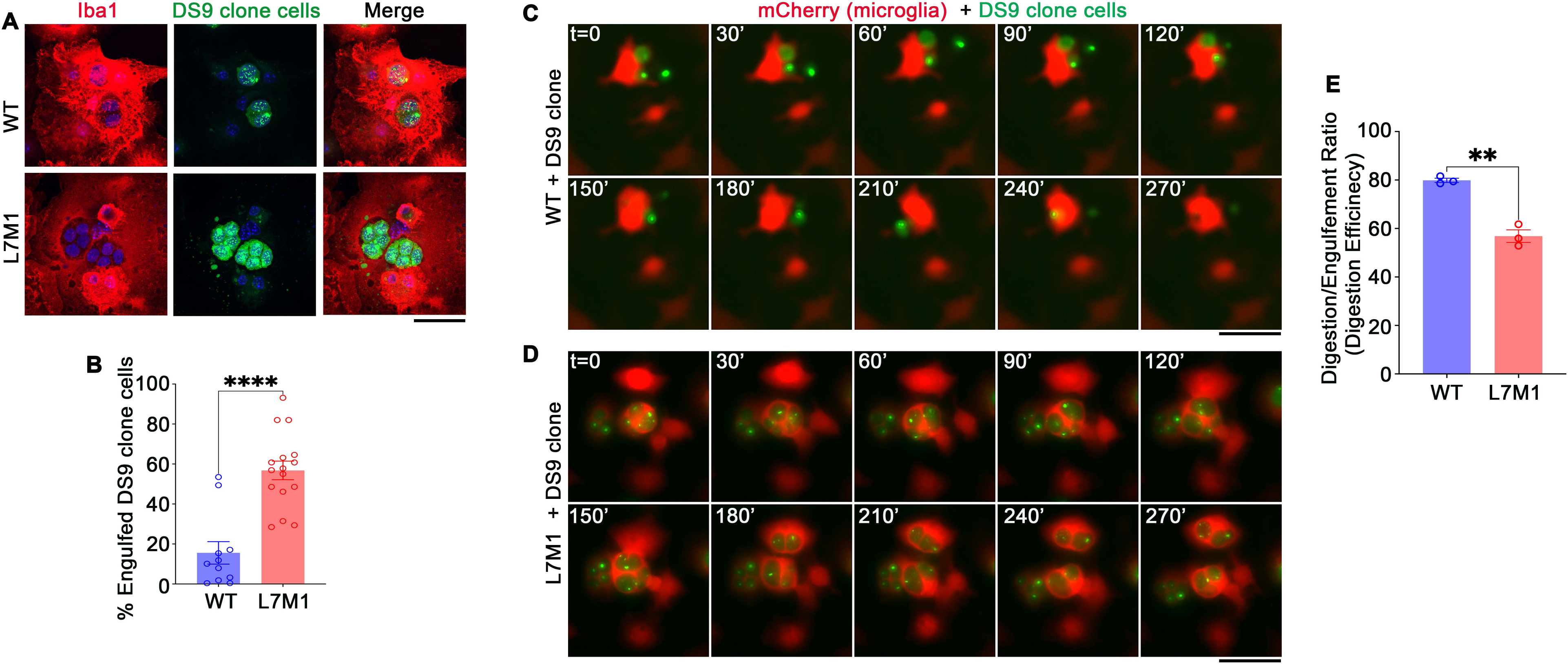
Immunoproteasome deficiency increases microglial cargo accumulation but impairs post-engulfment degradation. **(A)**Representative fluorescence images of WT and L7M1 primary microglia co-cultured with DS9 clone cells. Microglia were labeled with Iba1 (red), DS9 clone cell-derived GFP-tau signal is shown in green, and nuclei were counterstained with DAPI (blue). **(B)** Quantification of the percentage of DS9 clone cells engulfed by WT and L7M1 microglia. L7M1 microglia showed increased accumulation of DS9 clone cell material compared with WT microglia. Each point represents one independent experiment. In each experiment, two wells were analyzed, with three images acquired per well, for a total of six quantified images per experiment; values from the six images were averaged to generate one experimental replicate. WT, n = 11 experiments; L7M1, n = 16 experiments. **(C,D)** Representative frames from 22-h live-cell imaging movies showing WT **(C)** and L7M1 **(D)** mCherry-expressing microglia interacting with DS9 clone cell-derived GFP-positive tau cargo. Images were acquired every ∼15 min over 22 h; for display, every second acquired frame is shown at 30-min intervals over the first 270 min. Microglia are shown in red and DS9-derived tau cargo is shown in green. WT microglia progressively resolved internalized green cargo, whereas L7M1 microglia retained intracellular green cargo over time. Scale bars, 100 µm. **(E)** Quantification of digestion efficiency from the full 22-h live-imaging movies, expressed as the percentage of engulfed/engaged cargo that was subsequently digested or cleared: 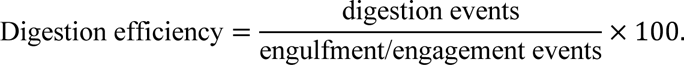 Engulfment/engagement was defined as GFP-positive DS9-derived cargo that contacted, overlapped with, or was internalized by red-labeled microglia. Digestion was defined as progressive disappearance or loss of detectable GFP-positive cargo after microglial uptake. Each plotted point represents one independent live-imaging experiment; two wells were analyzed per experiment and averaged to generate one experimental replicate. WT, n = 3 experiments; L7M1, n = 3 experiments. Data are shown as mean ± s.e.m. **P < 0.01, ****P < 0.0001 by two-tailed t-test. Scale bars for A, 20µm. Scale bars for C and D, 100µm.

### In vivo two-photon imaging reveals immunoproteasome-dependent clearance of engulfed tau by microglia

We next used longitudinal two-photon imaging to test whether immunoproteasome function is required for post-engulfment tau processing in the intact brain. CX3CR1^GFP/GFP^ reporter mice were crossed with L7M1 mice to generate homozygous L7M1/CX3CR1-GFP mice and CX3CR1-GFP littermate controls^30–32^ (n = 4 mice per group; **Fig. 5A**). At the time of cranial window implantation, AAV1-CBA-mCherry-TauP301L was delivered to the cortex to drive neuronal expression of human mutant TauP301L and mCherry^33–35^. Seven days later, repeated volumetric stacks were collected from the same fields at 0, 2, 4 and 6 h (**Fig. 5A**). Fluorescence contrast of large vascular structures enabled reliable longitudinal registration of the same imaging regions and cells across timepoints within individual mice. Three-dimensional rendering confirmed dense GFP-positive microglial surveillance near tau-P301L-mCherry-positive neurons (**Fig. 5B**), and orthogonal reconstructions verified that tau-mCherry signal was enclosed within microglial GFP volumes, supporting true engulfment rather than superficial overlap (**Fig. 5C**).

**Figure 5.**
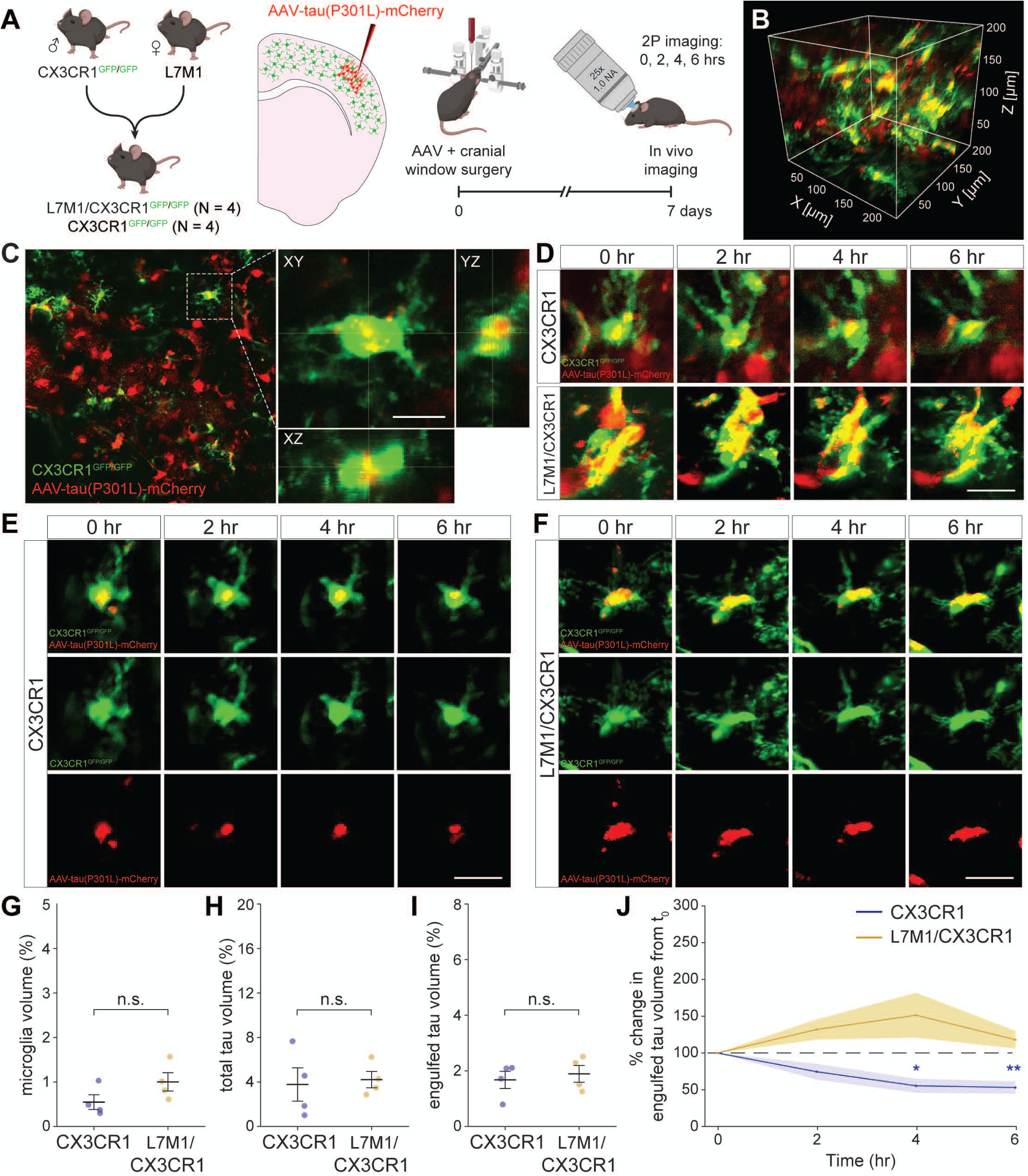
Immunoproteasome deficiency impairs in vivo resolution of microglia-engulfed tau. (**A**) Experimental design for longitudinal two-photon imaging. CX3CR1^GFP/GFP^ mice were crossed with L7M1 mice to generate CX3CR1^GFP/GFP^ controls and L7M1/CX3CR1^GFP/GFP^ immunoproteasome-deficient mice. Mice received AAV-tau(P301L)-mCherry together with cranial-window surgery, and microglia–tau interactions were imaged in vivo 7 days later at 0, 2, 4 and 6 h. CX3CR1-GFP n = 4 mice; L7M1/CX3CR1-GFP, n = 4 mice. (**B**) Representative three-dimensional two-photon reconstruction showing CX3CR1-GFP microglia in green and AAV-tau(P301L)-mCherry signal in red. (**C)** Representative two-photon image and orthogonal views showing tau-mCherry signal associated with GFP-positive microglia. (**D–F**) Representative longitudinal two-photon image sequences showing microglia-associated tau-mCherry signal over 6 h in CX3CR1-GFP and L7M1/CX3CR1-GFP mice. Red-channel images in **E** and **F** show the tau-mCherry signal used to track microglia-engulfed tau over time. Quantification at baseline of (**G**) microglial volume, (**H**) total tau-mCherry volume and (**I**) engulfed tau-mCherry volume showing no significant baseline differences between genotypes. (**J**) Longitudinal quantification of engulfed tau-mCherry volume expressed as percentage change from baseline. Engulfed tau decreased over time in CX3CR1-GFP microglia but was retained or increased in L7M1/CX3CR1-GFP microglia. Data are shown as mean ± s.e.m.; each point represents an individual quantified imaging region or object from n = 4 mice per genotype. n.s., not significant; **\***P < 0.05, **\*\***P < 0.01.

Longitudinal tracking revealed a genotype-dependent divergence in the fate of microglia-associated tau. In CX3CR1-GFP controls, tau-mCherry inclusions within microglia progressively diminished over 6 h (**Fig. 5D,E**). In L7M1/CX3CR1-GFP mice, microglia retained prominent tau-mCherry inclusions over time (**Fig. 5D,F**). Baseline microglial volume fraction, total tau-mCherry volume fraction and baseline engulfed tau volume fraction did not differ between genotypes (**Fig. 5G-I; n.s**.), indicating comparable microglial representation and tau availability at the start of imaging. Normalized longitudinal quantification showed that engulfed tau volume declined to approximately 50-60% of baseline by 4-6 h in control microglia, whereas L7M1/CX3CR1-GFP microglia exhibited an increase in engulfed tau volume, peaking at approximately 150% of baseline at 4 h and remaining above baseline at 6 h (**Fig. 5J; P < 0.05** at 4 h, **P < 0.01** at 6 h). These in vivo data identify an immunoproteasome-dependent step downstream of engulfment: microglia lacking inducible immunoproteasome catalytic capacity can engage and internalize tau, but fail to resolve that cargo efficiently on rapid time scales.

### Immunoproteasome deficiency drives microglial remodeling and phagolysosomal dysfunction

To define the transcriptional state associated with impaired cargo processing, we performed mouse single-nucleus RNA-seq in WT, APP/hTau dKI and L7M1/APP/hTau dKI mice. Feature plots, labeled UMAPs and marker-gene dot plots resolved the expected major CNS cell classes, including excitatory neurons, inhibitory neurons, microglia, astrocytes, oligodendrocytes, endothelial, ependymal cells and OPCs ^36–38^ (**Supplementary Figs. 3-5**). Because both the primary microglial co-culture assay and longitudinal in vivo two-photon imaging identified a defect in post-engulfment cargo resolution, we next asked whether immunoproteasome deficiency remodels microglial transcriptional programs involved in phagocytosis, phagosome maturation, and lysosomal degradation. Within microglia, APP/hTau dKI mice showed a disease-associated shift characterized by reduced homeostatic marker expression and increased activation-associated gene expression, including reduced P2ry12, Tmem119 and increased Trem2, Cst7 and Spp1^39–42^ relative to WT controls (**Fig. 6A,B**), whereas Aif1, Cx3cr1, Tyrobp and Apoe were not significantly altered in this comparison (**Fig. 6A,B**). Immunoproteasome deficiency further remodeled this disease-associated microglial state. Compared with APP/hTau dKI microglia, L7M1/APP/hTau dKI microglia showed further suppression of P2ry12, increased Trem2 expression and reduced expression of the cargo-processing genes Anxa3 and Mertk. However, other activation-associated genes, including Aif1, Tyrobp, Cst7, Spp1 and Apoe, were not further increased (**Fig. 6A,B**). These findings indicate that immunoproteasome deficiency does not simply amplify a broad activation program, but instead promotes a distinct P2ry12^low^/Trem2^high^ microglial state marked by loss of selected cargo-processing genes, most prominently Anxa3.

**Figure 6.**
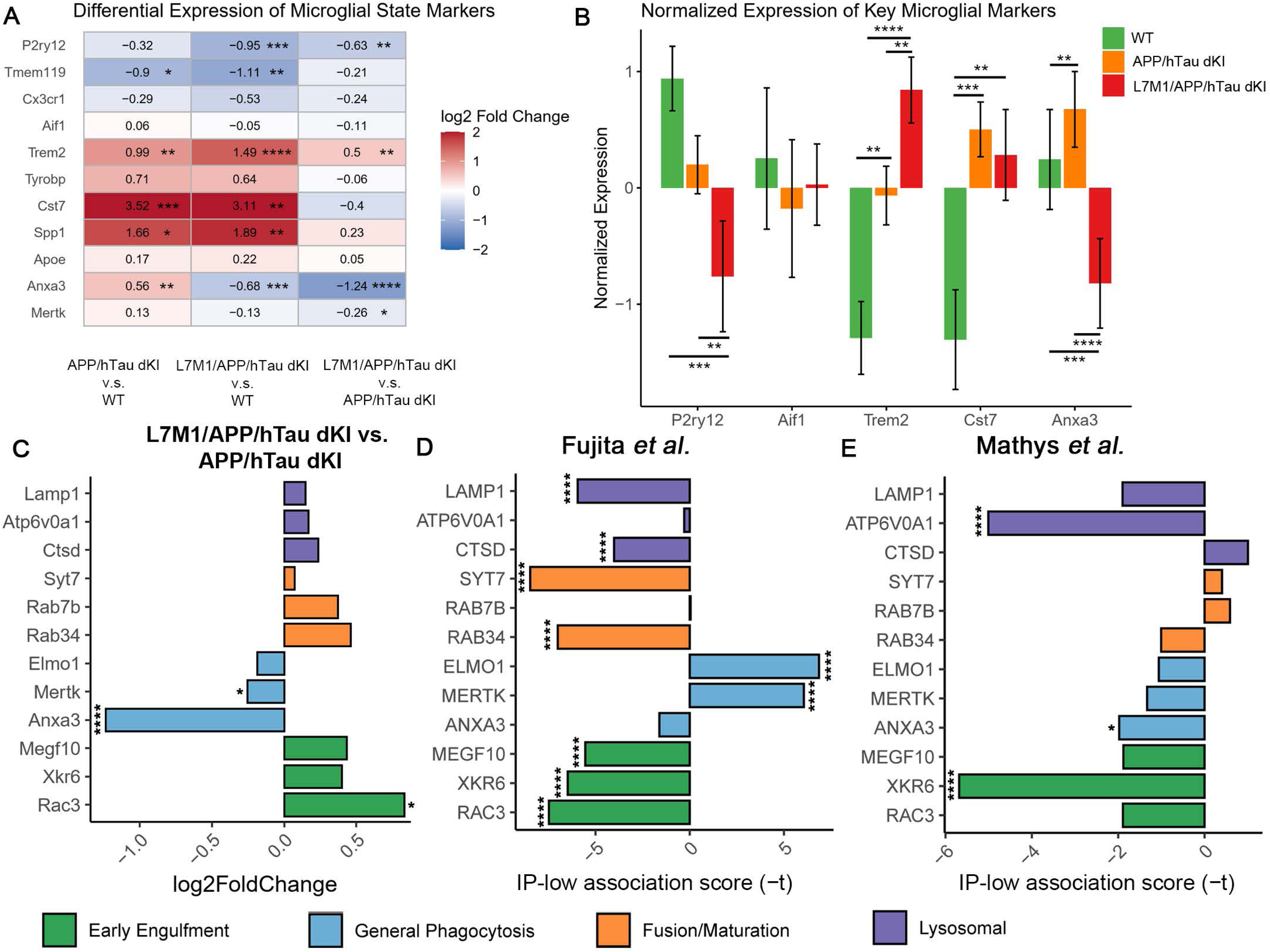
Immunoproteasome deficiency remodels microglial cargo-processing programs. **(A**)Heatmap showing differential expression of selected microglial state markers in mouse single-nucleus RNA-seq data across the indicated comparisons: APP/hTau dKI versus WT, L7M1/APP/hTau dKI versus WT and L7M1/APP/hTau dKI versus APP/hTau dKI. Values indicate log2 fold change, with red indicating increased expression and blue indicating reduced expression. (**B**) Normalized expression of selected microglial markers in WT, APP/hTau dKI and L7M1/APP/hTau dKI microglia, including the homeostatic marker P2ry12, the pan-microglial marker Aif1, disease-associated markers Trem2 and Cst7, and the cargo-processing gene Anxa3. (**C**) Differential expression of selected phagocytosis- and cargo-processing genes in L7M1/APP/hTau dKI microglia relative to APP/hTau dKI microglia. Genes are grouped by functional category: early engulfment, general phagocytosis, phagosome fusion/maturation and lysosomal degradation. Bars show log2 fold change. Reanalysis of human AD single-nucleus RNA-seq datasets from (**D**) Fujita *et al.* and (**E**) Mathys *et al.* showing covariate-adjusted associations between microglial immunoproteasome (IP) transcript expression and the same phagocytosis/cargo-processing genes shown in **C**. Bars show the negative partial-correlation *t* statistic, orienting the analysis toward the immunoproteasome-low state. Negative values indicate genes predicted to decrease in IP-low microglia, whereas positive values indicate genes predicted to increase in IP-low microglia. Corresponding expression scatter plots are shown in Supplementary Figs. 6 and 7; in those plots, gene expression is shown as a function of immunoproteasome expression, so positive scatter-plot slopes correspond to negative bars in D,E. Mouse snRNA-seq cohorts were sex balanced: WT, n = 4 mice; APP/hTau dKI, n = 6 mice; L7M1/APP/hTau dKI, n = 6 mice. Human single-nucleus RNA-seq sample sizes are provided in the Methods. Data are shown as mean ± s.e.m. where applicable. Statistical testing was performed as described in the Methods. ns, not significant; \**P* < 0.05, **P < 0.01, ***P < 0.001 and ****P < 0.0001.

We next examined a microglia-curated phagocytosis and phagolysosomal cargo-handling module spanning early cargo engagement (Megf10, Xkr6, Rac3), receptor/adaptor-mediated homeostatic phagocytosis (Elmo1, Mertk, Anxa3), phagosome maturation and membrane trafficking (Syt7, Rab7b, Rab34), and lysosomal fusion/degradation (Lamp1, Atp6v0a1, Ctsd)^43–46^ (**Fig. 6C**). Relative to APP/hTau dKI microglia, L7M1/APP/hTau dKI microglia showed increased expression of genes linked to early cargo engagement and actin-associated remodeling, including Megf10, Xkr6 and Rac3. Several trafficking and lysosomal genes, including Rab34, Rab7b, Lamp1, Atp6v0a1 and Ctsd, also showed upward trends. In contrast, Anxa3 was strongly reduced, Mertk was modestly but significantly reduced, and Elmo1 trended downward. Thus, immunoproteasome-deficient microglia retain transcriptional features of cargo engagement and lysosomal compartment activation, but show reduced expression of receptor/adaptor-linked cargo-processing genes. This split pattern is consistent with the imaging data, in which L7M1 microglia engage and engulf tau-bearing material but fail to efficiently complete intracellular cargo degradation.

To determine whether a related cargo-processing signature was present in human disease, we reanalyzed two independent ROSMAP human snRNA-seq datasets from Fujita et al.^47^ and Mathys et al.^48^ ROSMAP comprises longitudinal clinical-pathologic aging cohorts with cognitive assessments and postmortem neuropathologic characterization ^49,50^. The Fujita dataset included approximately 1.5 million transcriptomes from 424 aged individuals, whereas the Mathys dataset included approximately 2.3 million nuclei from 427 participants. After accounting for overlapping donors, these resources provided 619 non-overlapping brains and more than 3.9 million nuclei ^47,48^. For each dataset, analyses were restricted to annotated microglia. Donor-level microglial expression profiles were generated, and immunoproteasome abundance was modeled using variance-stabilized expression of PSMB8, PSMB9 and PSMB10. Associations between immunoproteasome expression and cargo-processing genes were then assessed using covariate-adjusted partial correlation (**Supplementary Figs. 6 and 7**). In Fig. 6D,E, the human analyses are displayed as the negative partial-correlation t statistic, so that the bars are oriented toward the immunoproteasome-low state. Under this convention, negative bars indicate genes predicted to decrease in IP-low microglia, whereas positive bars indicate genes predicted to increase in IP-low microglia.

In the Fujita dataset, lower immunoproteasome expression was associated with reduced expression of several genes involved in lysosomal degradation, phagosome maturation and early cargo handling, including LAMP1, CTSD, SYT7, RAB34, MEGF10, XKR6 and RAC3 (**Fig. 6D**). ANXA3 showed a weaker decrease, whereas RAB7B and ATP6V0A1 were near neutral. In contrast, ELMO1 and MERTK were increased in the IP-low state in this cohort.

In the Mathys dataset, IP-low microglia also showed reduced expression of multiple cargo-processing genes, including ATP6V0A1, RAB34, ELMO1, MERTK, ANXA3, MEGF10, XKR6 and RAC3 (**Fig. 6E**).

LAMP1, CTSD, SYT7 and RAB7B showed positive or near-neutral associations in this dataset. Although individual gene directions were not identical between cohorts, both human datasets supported remodeling of microglial cargo-processing programs in association with lower immunoproteasome expression. Notably, genes linked to early engulfment and actin-associated cargo handling, including MEGF10, XKR6 and RAC3, were recurrently decreased in the IP-low state across both cohorts.

The corresponding scatter plots in Supplementary Figs. 6 and 7 show the direct relationship between cargo-processing gene expression and immunoproteasome expression. Because Fig. 6D,E use the sign-inverted statistic, an upward slope in the supplementary scatter plots corresponds to a negative bar in Fig. 6D,E, indicating that the gene is predicted to be lower in the IP-low state. Conversely, a downward slope corresponds to a positive bar, indicating that the gene is predicted to be higher in the IP-low state.

Together, the mouse and human single-nucleus transcriptomic analyses support a model in which immunoproteasome deficiency, or low microglial immunoproteasome expression, does not prevent microglial activation or cargo engagement. Instead, it shifts microglia toward an activated but inefficient cargo-processing state, characterized by altered coordination of receptor/adaptor-mediated phagocytosis, phagosome maturation and lysosomal degradation.

### Low immunoproteasome states are associated with mitochondrial transcriptional suppression

Unbiased gene set enrichment analysis comparing L7M1/APP/hTau dKI mice with APP/hTau dKI controls revealed prominent negative enrichment of mitochondrial, ribosomal and protein homeostasis–related programs across CNS cell types (**Supplementary Fig. 8A**).

Negatively enriched pathways included mitochondrial membrane part, mitochondrial protein complex, mitochondrial matrix, NADH dehydrogenase complex assembly, NADH dehydrogenase complex, respiratory chain, oxidoreductase activity, cytochrome complex, mitochondrial gene expression, ribosome and translation (**Supplementary Fig. 8A**). This pattern was not consistent with generalized transcriptional collapse, as positively enriched pathways included extracellular matrix organization, cell adhesion, integrin-mediated signaling, cell-substrate junction, neuron projection guidance, semaphorin–plexin signaling, synapse organization, glutamatergic synapse, morphogenesis and broader tissue-remodeling programs (**Supplementary Fig. 8A**). Thus, immunoproteasome deficiency was associated with selective transcriptional reprogramming, with mitochondrial and protein homeostasis–related pathways emerging as one of the most consistent negatively enriched axes.

We therefore focused on mitochondrial pathway changes in Fig. 7. In the mouse snRNA-seq dataset, mitochondrial gene sets showed broad negative enrichment in L7M1/APP/hTau dKI mice relative to APP/hTau dKI controls across major CNS cell types, including excitatory neurons, inhibitory neurons, microglia, astrocytes, oligodendrocytes, endothelial cells, ependymal cells and OPCs (**Fig. 7A**). Negatively enriched mitochondrial terms included NADH dehydrogenase complex assembly, NADH dehydrogenase complex, mitochondrial protein complex, mitochondrial membrane part, mitochondrial matrix and mitochondrial gene expression.

**Figure 7.**
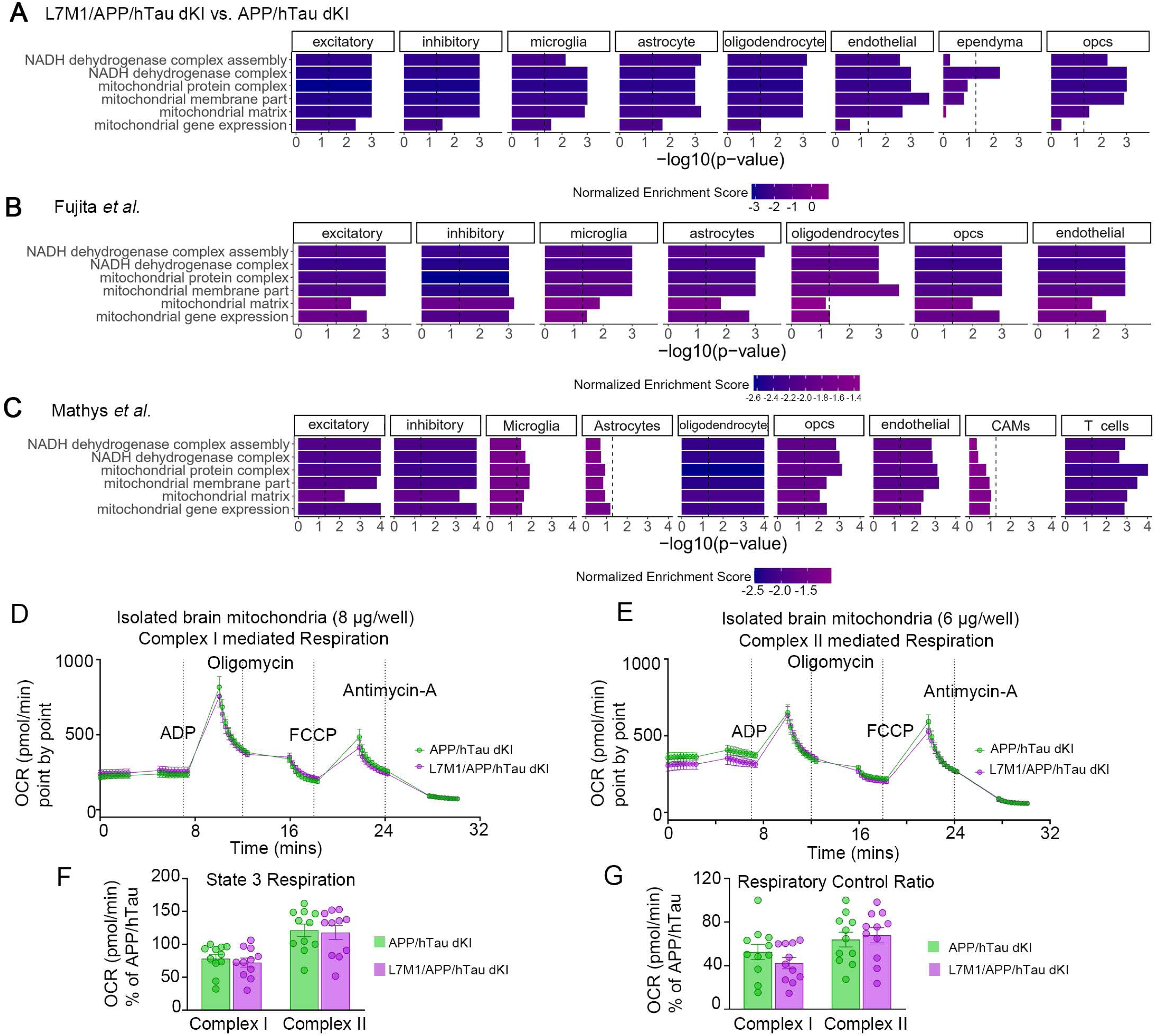
Immunoproteasome deficiency is associated with mitochondrial transcriptional suppression without detectable impairment of isolated brain mitochondrial respiration. Cell-type-resolved gene-set enrichment analysis of mitochondrial pathways in mouse and human single-nucleus RNA-seq datasets. (**A**) Single-nucleus RNA-seq datasets. Mouse snRNA-seq analysis comparing L7M1/APP/hTau dKI mice with APP/hTau dKI controls across excitatory neurons, inhibitory neurons, microglia, astrocytes, oligodendrocytes, endothelial cells, ependymal cells, and oligodendrocyte precursor cells. (**B**) Human snRNA-seq analysis from Fujita *et al.* showing mitochondrial gene-set enrichment associated with an immunoproteasome-low transcriptomic state across major CNS cell types. (**C**) Human snRNA-seq analysis from Mathys *et al.* showing mitochondrial gene-set enrichment associated with an immunoproteasome-low transcriptomic state across excitatory neurons, inhibitory neurons, microglia, astrocytes, oligodendrocytes, OPCs, endothelial cells, CNS-associated macrophages and T cells. Bars show −log10(*P* value), and bar color indicates normalized enrichment score. Negative normalized enrichment scores indicate reduced enrichment in the L7M1/APP/hTau dKI or human immunoproteasome-low state. Dashed vertical lines indicate the nominal significance threshold. Oxygen-consumption rate traces from isolated brain mitochondria from APP/hTau dKI and L7M1/APP/hTau dKI mice. (**D**) Complex I-mediated respiration measured using 8 µg mitochondria per well. (**E**) Complex II-mediated respiration measured using 6 µg mitochondria per well. ADP, oligomycin, FCCP and antimycin A were added at the indicated time points. Quantification of ADP-stimulated state 3 respiration (**F**) and respiratory control ratio (**G**) for complex I- and complex II-mediated respiration. No significant differences were detected between APP/hTau dKI and L7M1/APP/hTau dKI mitochondria under either respiratory condition. Data are shown as mean ± s.e.m.; each point represents an individual mitochondrial respiration measurement; n = 12 measurements per genotype for each condition. Statistical testing was performed as described in the Methods.

To determine whether a similar signature was present in human immunoproteasome-low states, we applied a continuous IP-low ranking strategy to the Fujita *et al.* and Mathys *et al.* human snRNA-seq datasets. For each cell type, genes were ranked using a signed IP-low association score derived from the relationship between normalized gene expression and the composite immunoproteasome score. The score was oriented toward the IP-low state by multiplying the association t-statistic by −1. Thus, genes that increased as immunoproteasome expression decreased received positive scores, whereas genes that decreased as immunoproteasome expression decreased received negative scores (**Supplementary Fig. 9**). For example, NT5C2 was negatively correlated with immunoproteasome expression and therefore received a positive IP-low GSEA input score, whereas IFI27L2 was positively correlated with immunoproteasome expression and therefore received a negative IP-low GSEA input score (**Supplementary Fig. 9**).

Using this IP-low scoring approach, human low-immunoproteasome states were again associated with negative enrichment of mitochondrial pathways across multiple CNS cell types in both datasets (**Fig. 7B,C; Supplementary Fig. 8B,C**). Negatively enriched pathways included mitochondrial gene expression, mitochondrial membrane part, mitochondrial protein complex, mitochondrial matrix, NADH dehydrogenase complex, NADH dehydrogenase complex assembly, cytochrome complex, ribosome and translation. These results support a conserved association between reduced immunoproteasome abundance and suppression of mitochondrial and protein homeostasis–related transcriptional programs.

At the individual-gene level, mitochondrial volcano plots further supported selective mitochondrial remodeling rather than uniform loss of all mitochondrial transcripts. In mouse snRNA-seq, L7M1/APP/hTau dKI mice showed broad remodeling of curated mitochondrial genes relative to APP/hTau dKI controls, with 315 genes decreased and 114 genes increased in the immunoproteasome-deficient disease state (**Supplementary Fig. 10A**). In the Fujita human snRNA-seq dataset, the IP-low state was associated with reduced expression of 404 mitochondrial genes and increased expression of 197 mitochondrial genes (**Supplementary Fig. 10B**). In the Mathys dataset, 192 mitochondrial genes were reduced and 409 mitochondrial genes were increased in the IP-low state (**Supplementary Fig. 10C**). Thus, although the pathway-level GSEA consistently identified negative enrichment of mitochondrial and proteostasis-related programs, individual mitochondrial transcripts showed dataset-specific bidirectional remodeling, consistent with altered mitochondrial state rather than generalized transcriptional failure.

We next tested whether this transcriptional mitochondrial signature was accompanied by overt impairment of bulk mitochondrial respiration. Isolated brain mitochondria from APP/hTau dKI and L7M1/APP/hTau dKI mice showed the expected oxygen-consumption responses to ADP, oligomycin, FCCP and antimycin A during complex I-supported respiration, with no detectable impairment in the immunoproteasome-deficient group (**Fig. 7D**). Complex II-supported respiration was similarly preserved (**Fig. 7E**). Quantification of ADP-stimulated state 3 respiration showed no significant reduction in either complex I-or complex II-mediated respiration in L7M1/APP/hTau dKI mitochondria compared with APP/hTau dKI controls (**Fig. 7F**). Respiratory control ratio was also maintained, indicating preserved mitochondrial coupling and respiratory responsiveness under these ex vivo assay conditions (**Fig. 7G**). Moreover, citrate synthase kinetic activity and normalized citrate synthase activity were not significantly different between groups, consistent with comparable mitochondrial enzymatic content in the assayed preparations (**Supplementary Fig. 11**).

Together, these data indicate that immunoproteasome deficiency and human IP-low states are associated with conserved suppression of mitochondrial and protein homeostasis–related transcriptional programs, but that isolated brain mitochondria from L7M1/APP/hTau dKI mice retain gross respiratory capacity. This divergence suggests that the transcriptional mitochondrial signature may reflect cell-state-specific metabolic stress, altered energetic allocation or local bioenergetic demand that is not captured by bulk isolated brain mitochondrial respiration assays.

## DISCUSSION

The role of immunoproteasome induction in AD has remained unresolved because this response can be interpreted either as an adaptive proteostatic mechanism or as a contributor to neuroinflammation. Immunoproteasomes were originally defined as cytokine-inducible proteasome variants that support antigen processing, but subsequent work established that they also help preserve protein homeostasis under oxidative and inflammatory stress^2,6,7^. In AD, increased immunoproteasome expression and activity in vulnerable regions and reactive glia have suggested a compensatory response to declining proteostasis, whereas studies using immunoproteasome inhibitors or single-subunit loss have supported a maladaptive inflammatory model by showing reduced cytokine output without major changes in amyloid deposition^9–12^. This ambiguity has left unresolved whether immunoproteasome induction in the AD brain primarily amplifies neuroinflammatory injury or preserves glial degradative capacity. Here, using genetic loss of inducible immunoproteasome catalytic subunits in complementary tau and amyloid–tau disease contexts, together with live cargo-handling assays, longitudinal in vivo two-photon imaging, mouse single-nucleus transcriptomics and human single-nucleus transcriptomic analyses, we identify the immunoproteasome as a protective component of glial proteostasis in AD-relevant brain environments.

Across both disease models, immunoproteasome deficiency increased pathological protein accumulation and heightened microglial reactivity without detectably suppressing constitutive 26S proteasome activity. This pattern points to a selective requirement for inducible immunoproteasome capacity under proteotoxic and inflammatory stress. Importantly, loss of immunoproteasome function did not dampen microglial activation, as would be expected if immunoproteasomes acted primarily as inflammatory drivers. Instead, microglia became more reactive in parallel with worsening tau and amyloid burden, suggesting that immunoproteasome deficiency promotes unresolved inflammatory engagement rather than immune quiescence.

The behavioral data are consistent with broader neuroimmune or circuit-level consequences of this state. L7M1-containing mice did not show a latency-based Morris water maze deficit, but they did show increased swim velocity, genotype-dependent increases in path length and greater peripheral occupancy. This pattern suggests a hyperkinetic or stress-responsive swim phenotype that complicates conventional interpretation of escape latency^25,26^. Because increased swim speed can mask inefficient navigation, preserved escape latency should not be interpreted as evidence of intact spatial cognition.

The central mechanistic insight of this study is that immunoproteasome deficiency impairs events that occur after pathological cargo has been engaged by microglia. In primary microglial co-cultures, L7M1 microglia accumulated more tau aggregate–bearing DS9-derived material than WT microglia. Live imaging showed that this accumulation reflected defective cargo resolution rather than more productive clearance: WT microglia progressively resolved internalized fluorescent cargo, whereas immunoproteasome-deficient microglia retained conspicuous intracellular material and showed a reduced digestion-to-engulfment ratio. Longitudinal two-photon imaging in the intact brain established the same principle in vivo. Baseline microglial volume, tau-mCherry availability and initial engulfed tau volume were comparable between genotypes, yet control microglia progressively reduced internalized tau cargo over hours, whereas L7M1 microglia retained or accumulated it. Thus, immunoproteasome deficiency does not prevent microglia from engaging or engulfing pathological material; instead, it uncouples engulfment from efficient intracellular cargo degradation.

This distinction is important for interpreting microglial responses in neurodegeneration. Microglia are often classified as protective or damaging based on aggregate association, uptake of pathological material or expression of activation markers^15,41,42^. Our data show that these readouts are insufficient to infer successful clearance. Immunoproteasome-deficient microglia remain cargo-engaged and reactive, but fail to complete the degradative process efficiently. We therefore propose that immunoproteasome deficiency drives a state of activation without resolution, in which microglia detect and internalize pathological material but cannot efficiently process it. Such a state would be expected to sustain intracellular stress, prolong inflammatory signaling and permit persistence or re-release of pathogenic protein species^51,52^. This model provides a mechanistic explanation for the coexistence of increased microgliosis and increased tau and amyloid burden in immunoproteasome-deficient disease models.

The transcriptional data support this model. In L7M1/APP/hTau dKI mice, microglia adopted a remodeled P2ry12l^ow^/Trem2^high^ with reduced expression of selected cargo-processing genes, most prominently Anxa3 and Mertk. At the same time, immunoproteasome-deficient microglia retained or increased components associated with cargo engagement, actin remodeling, trafficking and lysosomal compartment activation^39,41,42^. This split profile mirrors the imaging phenotype: microglia are not disengaged from pathological cargo, but their cargo-handling program is poorly coordinated. Increased lysosomal markers in this setting should therefore not be interpreted as evidence of successful degradation. Instead, they may reflect lysosomal expansion, stalled cargo processing or compensatory activation of an overloaded degradative pathway ^53–56^.

Human datasets provided an independent disease-relevant context for these findings. In two ROSMAP human single-nucleus RNA-seq cohorts, microglia with lower immunoproteasome transcript abundance showed pathway-level remodeling of the same phagocytosis-to-lysosome module altered in the L7M1/APP/hTau dKI model. Because the human analyses used continuous immunoproteasome expression rather than genetic deficiency, they should be interpreted as transcriptomic evidence of an immunoproteasome-low state rather than direct evidence of impaired immunoproteasome activity. Nevertheless, the convergence between mouse genetic data and human IP-low microglial signatures supports the idea that reduced immunoproteasome abundance is associated with altered cargo-processing programs in AD-relevant microglia.^36,53,57^.

The mitochondrial findings add a second layer to this framework. Immunoproteasome deficiency was associated with broad negative enrichment of mitochondrial respiratory-chain, mitochondrial gene-expression, ribosomal and protein homeostasis–related programs across mouse CNS cell types. Similar mitochondrial transcriptional signatures were observed in human IP-low states. However, isolated whole-brain mitochondria from L7M1/APP/hTau dKI mice retained complex I- and complex II-mediated respiration, respiratory control ratio and citrate synthase activity. This divergence suggests that the transcriptional mitochondrial signature does not reflect uniform failure of bulk mitochondrial respiration. Instead, it may indicate cell-state-specific metabolic stress, altered energetic allocation or localized bioenergetic demand that is not captured by whole-brain mitochondrial preparations. This possibility is especially relevant for microglia, because phagocytosis, phagosome maturation, lysosomal acidification and proteasome-dependent degradation are energetically demanding processes. Cell-type-resolved metabolic assays will be required to determine whether immunoproteasome deficiency creates local energetic bottlenecks at sites of cargo processing^58–61^.

Several limitations should be considered. The L7M1 model removes inducible immunoproteasome catalytic subunits broadly rather than selectively in microglia. Although the primary microglial assays, in vivo imaging and single-nucleus analyses strongly implicate microglial cargo-processing defects, contributions from astrocytes, neurons, endothelial cells, peripheral immune populations or developmental immune calibration cannot be excluded. In addition, PS19 and APP/hTau dKI mice model complementary aspects of tauopathy and amyloid–tau pathology but do not recapitulate the full complexity of sporadic human AD. The human single-nucleus transcriptomic analyses strengthen disease relevance but remain correlative and do not directly measure immunoproteasome activity, lysosomal flux or cargo degradation in human tissue. Finally, preserved bulk mitochondrial respiration does not rule out metabolic defects in specific cell types, disease-associated cell states or subcellular compartments.

Together, these findings shift the interpretation of immunoproteasome induction in AD. Rather than functioning primarily as a marker or amplifier of neuroinflammation, immunoproteasome engagement appears to support the degradative competence of disease-associated glia. When inducible immunoproteasome subunits are lost, microglia do not become inactive; they become reactive, cargo-loaded and inefficient. This failure of proteostatic resolution provides a unifying mechanism linking increased tau and amyloid burden, heightened microglial activation, altered behavioral state, disrupted phagolysosomal transcriptional programs and conserved human IP-low signatures.

In conclusion, the immunoproteasome supports efficient microglial processing of pathological tau and restrains amyloid and tau accumulation in AD-relevant brain environments. Immunoproteasome-deficient microglia can engage and engulf pathological material, but fail to clear it effectively, revealing a post-engulfment degradative checkpoint that depends on inducible proteasome capacity. These data position the immunoproteasome as a protective, stress-adaptive component of glial proteostasis and suggest that therapeutic strategies aimed at preserving or restoring immunoproteasome-dependent degradative competence may be preferable to broad suppression of immunoproteasome activity in AD.

## Materials and Methods

### Animals and experimental design

All animal procedures were approved by the Columbia University Irving Medical Center Institutional Animal Care and Use Committee and were performed in accordance with institutional guidelines and the U.S. National Institutes of Health Guide for the Care and Use of Laboratory Animals. Mice were maintained in a specific pathogen–free facility under controlled temperature and humidity conditions on a 12-h light/12-h dark cycle, with ad libitum access to standard chow and water unless otherwise stated. Animals were monitored routinely by veterinary staff and laboratory personnel. Both male and female mice were included, and sex was balanced across genotype groups whenever possible. Exact ages, sex distributions and sample sizes for each experiment are reported in the corresponding figure legends. Multiple mouse cohorts were used to assess the effects of impaired inducible immunoproteasome function across tauopathy, amyloid–tau pathology, in vivo microglial cargo handling, biochemical assays, mitochondrial assays, behavioral testing and primary microglial culture.

For the PS19 tauopathy experiments, mice included non-transgenic controls, L7M1 mice, PS19 mice and L7M1/PS19 mice. PS19 mice express human MAPT P301S under the murine prion promoter and are maintained on a mixed B6;C3 background B6;C3-Tg ((Prnp-MAPT*P301S)PS19Vle/J). And the L7M1 mice are lmp7−/−/mecl-1−/− on a C57BL/6-derived genetic background. Offspring that lacked the PS19 MAPT P301S transgene and were not immunoproteasome deficient were used as non-transgenic controls throughout the manuscript.

For amyloid–tau dKI experiments, APP/hTau dKI mice carry the App-NL-G-F-knock-in allele together with a humanized Mapt/hTau knock-in allele. The App-NL-G-F allele contains the familial AD Swedish, Arctic and Beyreuther/Iberian mutations, while the hTau knock-in allele replaces the endogenous mouse Mapt sequence with human MAPT sequence. These mice are knock-in models, from a C57BL/6-derived background. L7M1/APP/hTau dKI mice carried the same APP/hTau knock-in alleles and were additionally deficient for the inducible immunoproteasome subunits LMP7/β5i and MECL-1/β2i.

For longitudinal in vivo two-photon imaging of microglial tau handling, homozygous CX3CR1-GFP reporter mice and homozygous L7M1/CX3CR1-GFP mice were used. These mice expressed GFP from the endogenous CX3CR1 locus, enabling visualization of microglia in the intact brain. Thus, mice in the two-photon imaging cohort are referred to as CX3CR1-GFP reporter controls and L7M1/CX3CR1-GFP mice. For primary microglial experiments, microglia were isolated from postnatal day 1 C57BL/6 control (WT) and L7M1 pups.

### AAV-mediated tau expression and chronic cranial window implantation

Chronic cranial windows were implanted in 3- to 4-month-old homozygous CX3CR1-GFP mice and homozygous L7M1/CX3CR1-GFP mice for longitudinal in vivo two-photon imaging of microglial interactions with tau-expressing cortical tissue. The two-photon cohort included n = 4 mice per genotype group with an approximately 50:50 male:female distribution. Surgical procedures were performed under isoflurane anesthesia delivered in oxygen (5% induction, 2% maintenance). Body temperature was maintained at 36.6 °C using a feedback-controlled heating pad (FHC, Bowdoinham, ME). Perioperative analgesia included subcutaneous lidocaine at the incision site and buprenorphine (0.05 mg/kg), followed by ketoprofen (5 mg/kg) immediately after surgery and once daily for two additional postoperative days. Mice were secured in a stereotaxic frame, and the scalp was shaved and disinfected using alternating scrubs of antiseptic solution and 70% ethanol. A midline incision was made to expose the skull, and connective tissue was carefully removed. A thin layer of cyanoacrylate adhesive (Vetbond, 3M) was applied to the cleaned skull surface. A custom rectangular head bar was secured to the posterior skull using Metabond dental cement to permit repeated awake head fixation during longitudinal imaging. A 4-mm circular craniotomy was made over the primary somatosensory cortex (S1; anteroposterior = -1.5 mm, mediolateral = +3.2 mm relative to bregma). During drilling, sterile saline was applied intermittently to prevent thermal damage and to maintain tissue hydration. After the craniotomy was completed, AAV1-CBA-mCherry-TauP301L, titer >10^13^ GC/ml (www.vectorbuilder.com) was injected into the exposed cortex using pulled glass micropipettes (Drummond Scientific, Broomall, PA) connected to a Nanoliter 2020 precision injection system (World Precision Instruments, Sarasota, FL). A total volume of 300 nL was delivered at 5 nL/s at two dorsoventral depths (DV = -1.5 mm and -3.0 mm). After each injection, the micropipette was left in place for at least 2 min to minimize reflux and was then slowly withdrawn. Following viral delivery, the craniotomy was covered with a sterile glass coverslip and sealed with UV-curable dental cement to create a chronic optical window. Mice recovered on a warming pad and were monitored postoperatively according to institutional animal-care guidelines.

### In vivo two-photon imaging of microglial tau handling

One week after cranial window implantation and AAV1-CBA-mCherry-TauP301L injection, in vivo two-photon microscopy was performed to measure microglial engulfment and retention of tau-associated fluorescent cargo in the intact mouse cortex. Imaging was performed using an Ultima 2Plus two-photon microscope (Bruker, Madison, WI) equipped with an Insight X3 Dual tunable femtosecond laser (Spectra-Physics, Menlo Park, CA), non-descanned photomultiplier tube detectors (Hamamatsu Photonics, Hamamatsu, Japan) and a 25x water-immersion objective (1.0 NA; Nikon Instruments, Melville, NY).

The same cortical fields of view were re-identified across longitudinal imaging sessions using stable vascular landmarks visible through the cranial window. GFP-positive microglia and mCherry-positive tau signal were excited simultaneously at 980 nm. Laser power at the specimen was maintained below 20-30 mW to reduce phototoxicity during repeated imaging. Images were acquired at 1024 x 1024 pixels with a zoom factor of 1.5-2.0 and an approximate frame acquisition time of 5 s. Volumetric z-stacks were collected over 200-300 µm of cortical depth with a 2-µm z-step. Time-lapse volumetric imaging was performed by acquiring z-stacks every 2 h over a 6-h imaging period (0, 2, 4 and 6 h).

### Three-dimensional image processing and quantification of two-photon data

Two-photon image stacks were processed using custom ImageJ/Fiji macros and analyzed in three dimensions. For each mouse, raw stacks were cropped to the same cortical field of view across all timepoints. Residual motion and drift within and across imaging sessions were corrected before segmentation. Background signal was subtracted from both eGFP and mCherry channels using a rolling-ball algorithm with a 50-pixel radius. A 3D Gaussian filter (sigma = 1 voxel in x, y and z) was applied to reduce high-frequency noise while preserving cellular and punctate structures. For each mouse, image stacks from all timepoints were concatenated before thresholding so that segmentation parameters were constant across the entire longitudinal series. Global intensity means and standard deviations were calculated for each channel, and intensity thresholds were set at mean + 1 standard deviation for consistent within-animal segmentation. Binary masks were generated for GFP-positive microglia and mCherry-positive tau signal. Microglial masks were applied to corresponding tau masks using the Image Calculator function in ImageJ/Fiji to identify tau signal contained within microglial volumes. Tau-positive structures were quantified using the 3D Objects Counter plugin, and measurements including microglial volume, total tau volume, engulfed tau volume and non-engulfed tau volume were exported for downstream analysis in MATLAB and/or GraphPad Prism. Longitudinal change in microglia-associated tau was calculated by normalizing each timepoint to the baseline value at t = 0 h. Percent change in engulfed tau volume was calculated as ((engulfed tau volume at each timepoint - engulfed tau volume at baseline) / engulfed tau volume at baseline) x 100. For animal-level analyses, measurements from cells and fields belonging to the same mouse were averaged before statistical testing unless a nested or mixed-effects model was used.

### Native in-gel proteasome activity assay

Frozen mouse cortical tissue (50-100 mg) was homogenized on ice in native proteasome homogenization buffer containing 50 mM Tris-HCl (pH 7.4), 5 mM MgCl2, 5 mM ATP, 1 mM dithiothreitol (DTT), 1 mM EDTA, 10% glycerol, phosphatase inhibitor cocktail, 10 mM NaF and 25 mM beta-glycerophosphate. Homogenates were centrifuged at 14,000 x g for 25 min at 4 °C, and the soluble fraction was collected. Protein concentration was measured using the Pierce Bradford Plus Protein Assay Reagent (Thermo Fisher Scientific, 23238). Samples were normalized to 1 µg/µL, and filtered bromophenol blue was added at 4 µL per 200 µL sample to visualize migration without denaturing proteasome complexes. Equal protein amounts (20 µg per lane) were loaded onto native polyacrylamide 4 % gels and electrophoresed at 160 V for 4 h at 4 °C. Following electrophoresis, gels were incubated at 37 °C for 15min in assay buffer containing ATP, DTT and 150 µM Suc-LLVY-AMC (UBP Bio, G1101). Cleavage of Suc-LLVY-AMC was used as a readout of chymotrypsin-like proteasome activity. Fluorescent bands were imaged using an Azure Biosystems 600 imaging system. Band intensities corresponding to native proteasome activity were quantified in ImageJ/Fiji after background subtraction and were normalized as specified in the statistical analysis section.

### Western blotting

Mouse cortical tissue (50 mg) was homogenized on ice in RIPA lysis buffer supplemented with protease and phosphatase inhibitors. The final working composition of the RIPA buffer should be verified before submission. Homogenates were sonicated on ice at 50% amplitude for 10 s and centrifuged at 3,000 x g for 10 min at 4 °C. Supernatants were collected, and protein concentration was determined using the Pierce BCA Protein Assay Kit (Thermo Fisher Scientific, A55864). Protein lysates were normalized to 1 µg/µL in 4x NuPAGE LDS sample buffer (Invitrogen, NP0007) and denatured at 95 °C for 5 min. Equal protein amounts (20 µg per lane) were resolved on 4-12% Bis-Tris Midi gels (NuPAGE Bis-Tris Midi Protein Gels, 1.0 mm; Invitrogen, WG1402BOX and WG1403BOX) at 160 V until the dye front exited the gel. Proteins were transferred to 0.2-µm nitrocellulose membranes (Amersham Protran Premium, GE10600080) at 200 mA for 2 h. Membranes were blocked in 5% non-fat milk in TBST and incubated overnight at 4 °C with primary antibodies diluted in SuperBlock blocking buffer (Thermo Fisher Scientific, 37515). After primary antibody incubation, membranes were washed three times for 10 min each in TBST and incubated for 2 h at room temperature with HRP-conjugated anti-mouse or anti-rabbit secondary antibodies diluted in 5% milk/TBST. Chemiluminescent signal was detected using an Azure Biosystems 600 imaging system. Band intensities were quantified in ImageJ/Fiji after background subtraction. Target protein abundance was normalized to beta-actin or the indicated loading control and then expressed as fold change relative to the control group mean within the same blot or experimental batch.

### Immunohistochemistry and confocal microscopy

For histological analyses, mice were deeply anesthetized with a ketamine/xylazine cocktail until loss of pedal reflex and transcardially perfused with ice-cold 1× PBS to remove blood, followed by freshly prepared 4% paraformaldehyde (PFA) in PBS, pH 7.4. Brains were carefully removed and post-fixed in 4% PFA at 4°C for 24 h. After fixation, brains were rinsed in PBS and cryoprotected in 30% sucrose in PBS at 4°C until they sank. Cryoprotected brains were embedded in OCT, frozen on dry ice, and stored at −80°C until sectioning. Coronal brain sections were cut at 40 µm using a Cryostar NX50 cryostat. Sections were collected as free-floating sections and stored in PBS containing 0.02% sodium azide at 4°C until immunostaining. Anatomically matched sections were selected across animals. Quantified regions included cortex and hippocampus, including plaque-bearing cortical and hippocampal regions when applicable. Free-floating mouse brain coronal sections were transferred to 24-well plates and washed three times for 5 min each in PBST containing PBS and 0.3% Triton X-100. Sections were blocked for 1 h at room temperature in blocking buffer containing PBS, 0.3% Triton X-100 and 5% normal goat serum. Sections were then incubated overnight at 4°C with primary antibodies diluted in blocking buffer. Primary antibodies included markers for amyloid pathology, phospho-tau pathology, and microglial activation, including Aβ, phospho-tau, Iba1, and CD11b, are listed below. The following day, sections were washed three times for 10 min each in PBST and incubated for 2 h at room temperature with species-appropriate Alexa Fluor–conjugated secondary antibodies diluted 1:500 in blocking buffer. Secondary antibodies were selected according to the host species of the primary antibodies and included Alexa Fluor 488, Alexa Fluor 568 and Alexa Fluor 647 conjugates. Sections were protected from light during and after secondary antibody incubation. When used, nuclei were counterstained with DAPI. After secondary antibody incubation, sections were washed three additional times in PBS containing 0.02% Tween-20, mounted onto SuperFrost Plus glass slides, briefly air-dried and coverslipped using Fluoromount-G mounting medium. Slides were cured overnight at 4°C before imaging. Confocal images were acquired using a Zeiss LSM 800 laser-scanning confocal microscope with a 40× oil-immersion objective. Z-stacks were acquired through the tissue section in green, red and far-red channels. Within each staining experiment, identical acquisition parameters were used across experimental groups, including pinhole size, laser power, detector gain, z-step size and image resolution. Images were acquired from anatomically matched regions, and image acquisition and/or quantification were performed blinded to genotype whenever feasible. Scale bars represent 20 µm unless otherwise stated. Images were quantified using ImageJ/Fiji with constant thresholding, segmentation and region-of-interest definitions applied across all images within each experiment. Quantification was performed using anatomically matched sections and fields of view. For each mouse, four sections were analyzed, with three images acquired per section, yielding 12 images per mouse. The 12 image-level measurements were averaged to generate one biological replicate per animal; therefore, each plotted circle represents one mouse.

### Antibodies and key reagents

Primary and secondary antibodies used for Western blotting and immunohistochemistry are summarized below. Host species and catalog numbers should be verified before final submission, particularly for PHF1, AT8, Iba1, CD11b and 6E10.

| Application | Target/antibody | Vendor/catalog | Dilution and notes |
| --- | --- | --- | --- |
| Western blot | Beta-actin; mouse monoclonal | Cell Signaling Technology, 3700 | 1:10,000; loading control |
| Western blot | PHF1, pS396/pS404 tau; mouse | Gift from the late Dr. Peter Davies | 1:5,000 |
| Western blot | AT8, pS202/pT205 tau; mouse | Invitrogen, MN1020 | 1:1,000 |
| Western blot secondary | Anti-mouse IgG-HRP; goat anti-mouse | Jackson ImmunoResearch, 115-035-146 | 1:3,000 |
| Western blot secondary | Anti-rabbit IgG-HRP; donkey anti-rabbit | Invitrogen, 31458 | 1:3,000 |
| Immunohistochemistry | PHF1, pS396/pS404 tau; mouse | Gift from the late Dr. Peter Davies | 1:250 |
| Immunohistochemistry | AT8, pS202/pT205 tau; mouse | Invitrogen, MN1020 | 1:250 |
| Immunohistochemistry | Iba1; guinea pig | Synaptic Systems, 234 308 | 1:250 |
| Immunohistochemistry | CD11b | Invitrogen, 14-0112-82 | 1:100 |
| Immunohistochemistry | Aβ/6E10 | Novus Biologicals, NBP2-62566 | 1:250 |

### Morris water maze

Spatial learning and navigation were assessed in adult male and female mice with or without functional immunoproteasomes in the context of tau/amyloid pathology. Mice were tested over an 8-day Morris water maze protocol. During hidden-platform training, mice received four trials per day from randomized start positions. The hidden platform location remained fixed across all training trials and days. Each trial ended when the mouse reached the hidden platform or after a maximum of 60 s. Mice that did not locate the platform within 60 s were guided to the platform by the experimenter and allowed to remain on the platform for 5s before removal. Mouse movement was recorded and analyzed using EthoVision XT software (Noldus). Primary acquisition measures included latency to reach the hidden platform, swim velocity and total path length. Search paths and cumulative occupancy maps were generated from tracking data to evaluate search strategy, thigmotaxis/peripheral occupancy and spatial distribution during testing.

The PS19 cohort experiments used: Non-Tg n = 36; L7M1 n = 38; PS19 n = 25; L7M1/PS19 n = 25, and the experiments were done in two separate studies due to the large number of animals. The APP/hTau dKI cohort experiment used WT n = 14; L7M1 n = 11; APP/hTau dKI n = 13; L7M1/APP/hTau dKI n = 13.

### Primary microglial culture and live-cell imaging of tau-seed uptake

Primary microglia were isolated from mixed glial cultures prepared from postnatal day 1 WT or L7M1 mouse pups. Microglia were plated at 60,000 cells per well in 12-well plates coated with 0.1 mg/mL poly-D-lysine (Gibco, A3890401). After approximately two weeks of culturing, microglia were transduced with pre-packaged lentiviral particles expressing mCherry fluorescent protein (rLV.EF1.mCherry-9; Takara Bio, 0037VCT) at a multiplicity of infection. The medium was changed the following day, and mCherry expression was allowed to develop for 7 days. On day 6 after lentiviral transduction, 40,000 DS9 cells were added to each well and co-cultured with primary microglia for 48 h. DS9 cells are a clonal HEK293-derived line that stably propagates tau aggregates and releases misfolded YFP-tagged tau into the culture medium, thereby providing seed-competent tau cargo for microglial uptake. Live-cell imaging was performed for 24 h using a BioTek Cytation 5 Cell Imaging Multi-Mode Reader with BioSpa environmental control (Agilent). Images were acquired every 15 min for 22 h at 20x magnification in the Texas Red/mCherry and GFP/YFP fluorescence channels. Time-lapse image stacks from each fluorescence channel were aligned and merged in ImageJ/Fiji. Merged image sequences were exported as .avi files for visualization. For quantitative analyses, mCherry-positive microglia and YFP-positive tau signal were segmented using identical thresholds across experimental groups. Readouts included total YFP area or volume within microglial masks, number of YFP-positive puncta, integrated YFP intensity, fraction of YFP-positive microglia and time-dependent loss or retention of internalized YFP signal. Well-level or independent-culture-level averages were used for statistical testing to avoid treating repeated images or cells from the same culture as independent biological replicates.

### Isolation of enriched mitochondrial fractions from the mouse cortex

Fresh enriched mitochondrial fractions were prepared from the APP/hTau dKI and L7M1/APP/hTau dKI mice cortex for extracellular flux analysis. Age-matched mice, approximately 10-12 months of age, were euthanized by cervical dislocation on the day of the assay. Brains were rapidly removed and placed in cold 1x PBS. Cortical tissue was dissected, washed three times in cold PBS to remove blood and cut into small pieces on ice in MSHE homogenization buffer containing 210 mM mannitol, 70 mM sucrose, 5 mM HEPES and 1 mM EGTA, adjusted to pH 7.2 with KOH. Tissue was homogenized with approximately 4-5 strokes using a Teflon Dounce homogenizer. The homogenate was diluted twofold with MSHE wash buffer containing 210 mM mannitol, 70 mM sucrose, 5 mM HEPES, 1 mM EGTA and 0.5% fatty-acid-free BSA, pH 7.2. Homogenates were centrifuged at 900 x g for 10 min at 4 °C, and the supernatant was collected and subjected to a second 900 x g spin for 10 min at 4 °C to remove remaining debris and nuclei. The resulting supernatant was centrifuged at 9,000 x g for 10 min at 4 °C. The mitochondrial pellet was resuspended in 1.5 mL MSHE homogenization buffer and centrifuged at 8,000 x g for 10 min at 4 °C. The final enriched crude mitochondrial pellet was resuspended in 100 µL MSHE homogenization buffer and kept on ice until loading into Seahorse assay plates. Protein concentration of the mitochondrial preparation was measured using the Pierce Bradford Plus Protein Assay Reagent (Thermo Fisher Scientific, 23238) on a Tecan Spark microplate reader. All steps were performed on ice or at 4 °C to preserve mitochondrial function.

### Seahorse extracellular flux analysis of isolated cortical mitochondria

Mitochondrial oxygen consumption rate (OCR) was measured using an Agilent Seahorse XFe96 analyzer. Seahorse sensor cartridges (Agilent XFe96/XF Pro FluxPak, 103792-100) were hydrated the day before the assay according to the manufacturer’s instructions. Immediately before loading, enriched mitochondrial fractions were diluted in mitochondrial assay solution containing 220 mM mannitol, 70 mM sucrose, 5 mM KH2PO4, 5 mM MgCl2·6H2O, 2 mM HEPES, 1 mM EGTA and 0.1% fatty-acid-free BSA, adjusted to pH 7.2 with KOH. Complex I-dependent respiration was measured in assay solution supplemented with 5 mM pyruvate and 5 mM malate. Complex II-dependent respiration was measured in assay solution supplemented with 5 mM succinate and 2 µM rotenone. Mitochondria were plated at 8 µg protein per well for complex I assays and 6 µg protein per well for complex II assays in a 50-µL loading volume. Plates were centrifuged at 2,000 x g for 10 min at 4 °C to adhere mitochondria to the well bottom, after which 130 µL additional assay buffer was added to each well. Background wells contained assay buffer without mitochondria. Injection ports were loaded with ADP, oligomycin, FCCP and antimycin A diluted in the corresponding substrate-containing assay buffer. Final assay concentrations after injection were 4 mM ADP, 4 µM oligomycin, 6 µM FCCP and 4 µM antimycin A. OCR was measured at basal substrate-supported respiration and after sequential injection of ADP, oligomycin, FCCP and antimycin A. Respiratory-state readouts included substrate-supported state 2 respiration, ADP-stimulated state 3 respiration, oligomycin-insensitive state 4o respiration, FCCP-stimulated maximal respiration and antimycin A-insensitive background respiration. Background OCR was subtracted before downstream analysis.

### Citrate synthase activity assay

Citrate synthase activity was measured in mouse cortical tissue from age-matched APP/hTau dKI and L7M1/APP/hTau dKI mice approximately 10 months of age. Mice were euthanized by cervical dislocation on the day of the assay, and brains were rapidly harvested into cold 1x PBS. Cortical tissue was washed three times in cold PBS to remove residual blood. Approximately 10 mg cortex was used for the citrate synthase assay, and remaining tissue was snap-frozen for additional analyses. Cortical tissue was homogenized in 100 µL assay buffer according to the manufacturer’s protocol (Abcam citrate synthase assay kit, ab239712). Homogenates were centrifuged at 10,000 rpm for 5 min at 4 °C, and supernatants were collected for the assay. Reagents were equilibrated to room temperature before use, while samples were maintained on ice. A reduced-glutathione standard curve was prepared by dilution of the kit-provided 20 mM GSH standard to generate 0, 8, 16, 24, 32 and 48 nmol/well standards. Samples were diluted 25-fold; standards, samples and kit-provided positive controls were added to a 96-well plate in 50-µL volumes. Reaction master mix containing assay buffer, DTNB and citrate synthase substrate was added to sample and standard wells. Background-control wells received assay buffer and DTNB without citrate synthase substrate. Absorbance at 412 nm was recorded kinetically for 40 min at 25 °C. Protein concentration was determined by Bradford assay. Citrate synthase activity was calculated from the linear increase in absorbance using the standard curve and normalized to protein concentration. Activity was calculated as B / (deltaT x V) x D, where B is the amount of CoA-SH generated between selected timepoints, deltaT is the reaction time interval, V is the sample volume and D is the sample dilution factor.

### Mouse brain nuclei isolation for snRNA-seq

Single-nucleus RNA-seq was performed using nuclei isolated from brain tissue of age- and sex-matched WT, APP/hTau dKI, and L7M1/APP/hTau dKI mice. Nuclei were isolated using an optimized detergent-based protocol adapted from the DroNc-seq nuclei isolation approach described by Habib et al^1^ for downstream 10x Genomics single-nucleus capture. All steps were performed on ice or at 4°C using pre-chilled buffers, tubes, filters, pestles and centrifuge rotors to preserve nuclear integrity and RNA quality. Frozen hippocampal slices were placed in a 1.5-mL microcentrifuge tube containing 50 µL ice-cold Nuclei EZ lysis buffer supplemented with RNase inhibitor and mechanically dissociated with a disposable pestle. An additional 25 µL lysis buffer was used to rinse residual tissue from the pestle, and samples were kept on ice for a total lysis time of 5 min. Samples were centrifuged at 500 × g for 5 min at 4°C, and the nuclear pellet was gently resuspended in 50 µL ice-cold Nuclei EZ lysis buffer using wide-bore tips and incubated on ice for another 5 min. PBS containing RNase inhibitor and 0.04% BSA was then added, and nuclei were pelleted again at 500 × g for 5 min at 4°C. The pellet was resuspended in 100 µL PBS/0.04% BSA and passed through a pre-wetted 40-µm pluriStrainer to remove debris and aggregates. Nuclei were inspected and counted by Trypan Blue staining, centrifuged again at 500 × g for 5 min at 4°C, resuspended in 100 µL PBS/0.04% BSA, and passed through a pre-wetted 5-µm filter. Final nuclei suspensions were re-counted, inspected microscopically for integrity and clumping, diluted to the concentration required by the Columbia Genome Center at CUIMC and submitted immediately on ice for 10x Genomics Chromium single-nucleus capture, library preparation and sequencing. The Core performed encapsulation, library preparation, sequencing and primary processing, while downstream quality control, clustering, annotation, pseudobulk aggregation and statistical analyses were performed by our group.

### Mouse snRNA-seq downstream quality control, normalization, clustering and annotation

Core-generated raw or filtered count matrices were imported into R and processed using Seurat 5.3.0. Quality-control metrics were calculated for each nucleus, including total UMIs, number of detected genes, mitochondrial read fraction, ribosomal read fraction, sample identity and library identity. Sample-level QC distributions were reviewed before filtering to ensure that thresholds removed technical outliers without eliminating biologically relevant nuclei. Low-quality nuclei and likely multiplets were removed before clustering. Nuclei with fewer than 200 detected genes, more than 7,000 detected genes, fewer than 500 UMIs or more than 25,000 UMIs were excluded. Additional doublets and multiplets were identified using DoubletFinder 2.0 with expected doublet rates based on recovered nuclei per library. Doublet calls were inspected together with UMAP location and co-expression of mutually exclusive cell-type markers, and doublet-assigned nuclei were excluded from downstream analyses. Highly variable genes were identified, and expression values were scaled before principal-component analysis. Principal components used for graph construction were selected based on variance explained, elbow plots and preservation of major biological cell classes. Retained principal components were taken as the input for UMAP for visualization, and nuclei were clustered using a graph-based Louvain or Leiden algorithm. Clusters were annotated using canonical CNS marker genes and manual review of marker expression. Excitatory neurons were identified by markers such as Slc17a7, Camk2a, Neurod6 and Nrgn; inhibitory neurons by Gad1, Gad2 and Gabbr1; microglia by Csf1r, Cx3cr1, P2ry12, Hexb and Tmem119; astrocytes by Aqp4, Fgfr3, Slc1a2 and Gja1; oligodendrocytes by Mag, Mog, Mbp and Plp1; oligodendrocyte precursor cells by Pdgfra, Cspg4 and Vcan; endothelial cells by Flt1, Kdr, Cldn5 and Ly6c1; pericytes and vascular mural cells by Pdgfrb, Rgs5 and Acta2; and ependymal cells by Foxj1 and Dnah11. Clusters dominated by low-quality nuclei, mixed-lineage markers or doublet-like profiles were excluded or conservatively annotated as ambiguous. Microglia were subset for focused analysis. Within the microglial subset, counts were re-normalized, variable genes were recalculated and clustering was repeated to resolve microglial states. Microglial states were interpreted using homeostatic markers such as P2ry12, Tmem119, Hexb and Sall1; disease-associated and Trem2-dependent markers such as Trem2, Apoe, Tyrobp, Lpl and Cst7; inflammatory markers; phagocytic and cargo-handling genes; phagosome maturation genes; lysosomal genes; and proteasome/immunoproteasome genes. For biological replicate-aware analyses, raw counts were pseudobulk-aggregated after QC by summing nucleus-level counts for each gene within each mouse and cell type. For microglia-focused analyses, counts were also pseudobulk-aggregated by mouse within microglial state or subcluster when sample size permitted. Pseudobulk aggregation was used to preserve the animal as the biological replicate and to avoid treating individual nuclei as independent samples.

### Human ROSMAP snRNA-seq datasets and preprocessing

Human AD snRNA-seq datasets were obtained from the AMP-AD Knowledge Portal, including the Mathys *et al*. ROSMAP dataset^2^ (syn52293417) and the Fujita *et al*. ROSMAP dataset ^3^(syn51123521). Processed count matrices, cell-type annotations and donor metadata were used when available. When raw count matrices were used, the data were processed using a pipeline matched as closely as possible to the mouse snRNA-seq analysis. Analyses were performed separately for each human cohort and then compared at the level of cell types, genes and pathways. Nuclei were grouped by donor and cell type, and raw counts were summed by gene within each donor-cell type combination to generate pseudobulk count matrices. Donor-level covariates were extracted from available metadata, including sex, age at death, postmortem interval, race, education, diagnosis, Braak stage, *APOE* genotype and sequencing/library metrics, as available for each cohort. Pseudobulk count matrices were variance-stabilizing normalized using DESeq2 for downstream modeling and visualization. Nuclear-encoded mitochondrial genes were analyzed using a strict curated mitochondrial reference list^4^. Mis-aligned mitochondrial genome-encoded MT-* genes due to sequence homogeneity were excluded from nuclear mitochondrial module analyses and cross-species mitochondrial comparisons because snRNA-seq is designed to capture nuclear RNA and abundant mitochondrial transcripts can reflect residual cytoplasmic or mitochondrial carryover rather than nuclear transcriptional regulation.

### Statistical analyses

#### General statistical framework for non-transcriptomic experiments

Statistical analyses for non-transcriptomic experiments were performed using GraphPad Prism, version 10.1.0, unless otherwise stated. Image processing and densitometry were performed in ImageJ/Fiji, with exported values organized in GraphPad Prism, MATLAB and/or Microsoft Excel for visualization and statistical analysis. All statistical tests were two-sided. Data are presented as mean ± SEM unless otherwise stated in the figure legends.

For in vivo mouse experiments, the individual mouse was considered the biological replicate. For cell-culture assays, the independent primary microglial preparation, pup, litter or culture batch was considered the biological replicate; repeated wells, images or cells from the same preparation were treated as technical measurements. Technical replicates were averaged before group-level statistical testing. Normality was assessed using Shapiro-Wilk tests and inspection of residuals when sample size permitted. For comparisons between two groups, normally distributed data were analyzed using unpaired two-tailed Student’s t tests; Welch’s correction was used when variances were unequal. Non-normally distributed two-group comparisons were analyzed using Mann-Whitney U tests. For comparisons involving more than two groups or more than one factor, one-way ANOVA, two-way ANOVA, repeated-measures ANOVA or mixed-effects models were used as appropriate. Multiple comparisons following ANOVA were corrected using Tukey’s post hoc tests as specified in the relevant figure legend. P < 0.05 was considered statistically significant unless a false-discovery-rate threshold or other correction was specified.

#### Statistical analysis of in vivo two-photon imaging

For longitudinal in vivo two-photon imaging, the primary outcome was the change in engulfed tau volume within eGFP-positive microglia over time. Cell-level values were first normalized to the baseline timepoint for each cell or field of view. To avoid pseudoreplication, values from multiple microglia or fields of view within the same mouse were averaged to generate one value per mouse per timepoint before Prism-based group comparisons. Genotype and time were analyzed using two-way repeated-measures ANOVA with time as the within-subject factor and genotype as the between-subject factor. When any timepoint was missing, the equivalent Prism mixed-effects model with restricted maximum likelihood was used. Post hoc comparisons between genotypes at individual timepoints were corrected using Sidak’s multiple-comparisons test. Baseline measures, including initial microglial volume, initial tau-mCherry signal, baseline engulfed tau volume and baseline non-engulfed tau volume, were compared between genotypes using unpaired two-tailed t tests or Mann-Whitney U tests depending on distribution. If single-cell values are displayed in the figures, each cell should be shown as a nested datapoint, but statistical inference should be performed using animal-level averages or a nested/mixed-effects model with mouse identity included as a random or nested factor.

#### Statistical analysis of native proteasome activity and Western blotting

Native in-gel proteasome activity was quantified from background-subtracted band intensities. When bands corresponding to distinct proteasome complexes were resolved, each band was analyzed separately. Intensities were normalized to the control group mean or to total lane signal when appropriate and plotted as fold change relative to control. Two-group comparisons were analyzed using unpaired two-tailed t tests or Mann-Whitney U tests. Western blot target-band intensities were normalized to beta-actin or the indicated loading control within the same lane. Normalized values were then expressed as fold change relative to the mean of the control group within the same blot or experimental batch. Two-group comparisons were analyzed using unpaired two-tailed t tests with Welch’s correction when appropriate.

#### Statistical analysis of immunohistochemistry

Confocal images were quantified using identical acquisition and thresholding parameters across groups. For each marker and brain region, field-level or section-level values from the same animal were averaged to generate one biological replicate per mouse. Quantified histological outcomes included, as applicable, percent area covered by immunoreactivity, integrated fluorescence intensity, number or density of positive objects, plaque burden, phospho-tau burden, microglial marker burden and microglial morphological or colocalization metrics. Two-group comparisons were performed using unpaired two-tailed t tests or Mann-Whitney U tests.

#### Statistical analysis of Morris water maze

Morris water maze acquisition measures, including escape latency, swim velocity and path length, were analyzed in GraphPad Prism using two-way repeated-measures ANOVA with training day as the repeated within-subject factor and genotype or experimental group as the between-subject factor. Bonferroni multiple-comparisons test was used for genotype comparisons within individual training days. Because swim speed can influence escape latency, latency results were interpreted together with swim velocity, path length and search-strategy/occupancy metrics.

#### Statistical analysis of live-cell tau uptake assays

Live-cell tau uptake assays were analyzed from time-lapse images of primary WT and immunoproteasome-deficient microglia co-cultured with DS9 tau-seeding cells. Identical imaging and segmentation parameters were applied across groups. Microglial area was defined by mCherry fluorescence, and DS9-derived tau cargo was quantified from the YFP/GFP-positive signal. Engulfment events were defined as internalization of tau-positive material by mCherry-positive microglia, whereas digestion/clearance events were defined as reduction or disappearance of internalized tau signal over time. For each independent culture, replicate wells, fields or movies were averaged before statistical comparison. Engulfment was normalized to microglial area and/or microglial number, and digestion efficiency was calculated as the ratio of clearance events to engulfment events. Endpoint comparisons between WT and immunoproteasome-deficient microglia, including engulfment rate, retained tau burden and digestion-to-engulfment ratio, were analyzed using unpaired two-tailed t tests

#### Statistical analysis of Seahorse mitochondrial respiration

Seahorse OCR values were background-subtracted using wells containing assay buffer without mitochondria. Technical wells for each mouse, substrate condition and injection sequence were averaged before statistical analysis, and the individual mouse or mitochondrial preparation was treated as the biological replicate. Complex I- and complex II-dependent respiration were analyzed separately. Respiratory parameters included state 2 respiration, ADP-stimulated state 3 respiration, oligomycin-insensitive state 4o respiration, FCCP-stimulated maximal respiration, antimycin A-insensitive residual OCR, respiratory control ratio and other derived indices where indicated. Respiratory control ratio was calculated as ADP-stimulated state 3 OCR divided by oligomycin-insensitive state 4o OCR unless otherwise specified. OCR traces across injection steps were analyzed using two-way repeated-measures ANOVA or mixed-effects analysis with injection state as the repeated factor and genotype as the between-group factor. Individual derived respiratory parameters were compared between genotypes using unpaired two-tailed t tests or Mann-Whitney U tests. For experiments including more than two genotypes or conditions, one-way or two-way ANOVA was used with correction for multiple comparisons. Wells with failed injection, unstable baseline, loss of mitochondrial adherence or values outside predefined technical QC thresholds were excluded before averaging.

#### Statistical analysis of citrate synthase activity

Citrate synthase activity was calculated from the linear portion of the kinetic absorbance curve at 412 nm and normalized to protein concentration for each sample. Technical replicate wells were averaged to generate one value per mouse. APP/hTau dKI and L7M1/APP/hTau dKI groups were compared using unpaired two-tailed t tests when assumptions of normality and equal variance were met; otherwise, Welch’s t test or Mann-Whitney U test was used. Data were plotted as activity normalized to protein concentration and, where appropriate, as fold change relative to APP/hTau double knock-in controls.

#### General statistical framework for transcriptomic and spatial datasets

Transcriptomic statistical analyses were designed to preserve the biological replicate structure of each dataset. For mouse snRNA-seq, the animal was treated as the biological replicate. For human snRNA-seq and spatial transcriptomic datasets, the donor or sample was treated as the biological replicate or repeated-measures unit. Individual nuclei, cells or spatial units were not treated as independent biological replicates when animal- or donor-level replication was available. Unless otherwise stated, multiple testing was controlled using the Benjamini-Hochberg false-discovery rate (FDR) procedure.

#### Mouse pseudobulk differential-expression analysis

For each major cell type, pseudobulk raw count matrices were analyzed using DESeq2 1.52.0. Genes with low counts across pseudobulk samples were filtered before modeling. The primary mouse model compared L7M1/APP/hTau double knock-in mice with APP/hTau double knock-in controls and included sex as a covariate: counts ∼ sex + genotype When supported by sample size and study design, additional covariates such as sequencing batch, library preparation batch or age were included or assessed in sensitivity analyses. DESeq2 Wald statistics were used to rank genes for preranked gene-set enrichment analysis. For visualization, heatmaps, principal-component analysis and module summaries, pseudobulk counts were transformed using the DESeq2 variance-stabilizing transformation. Differentially expressed genes were defined using FDR correction (FDR < 0.05) with effect-size thresholds reported where applied.

#### Human immunoproteasome-low regression modeling

Human datasets were used to model transcriptional consequences of reduced immunoproteasome expression as an analogue of the mouse immunoproteasome-deficient state. Immunoproteasome expression was defined a priori as the mean variance stabilizing-transformed of *PSMB8*, *PSMB9* and *PSMB10*. Models were performed within each cell type, with emphasis on microglia.

For the Mathys *et al.* ROSMAP dataset, each gene was modeled as:

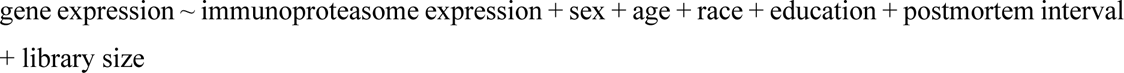

For the Fujita *et al*. ROSMAP dataset, each gene was modeled as:

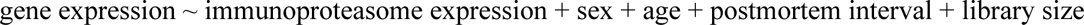

Library size refers to the donor-level sequencing-depth or DESeq2-derived normalization metric used to account for residual depth effects when variance-stabilized expression values were modeled directly. When available and appropriate, diagnosis, Braak stage, *APOE* genotype and sequencing batch were assessed in sensitivity models to ensure that immunoproteasome-associated signatures were not driven solely by disease severity or technical variation.

The regression coefficient for immunoproteasome expression reflects the association of each gene with higher immunoproteasome expression. To represent the predicted direction under immunoproteasome reduction, the *t* statistic derived from the coefficient for immunoproteasome expression was multiplied by -1. Thus, a positive immunoproteasome-deficiency direction statistic indicates genes predicted to increase when immunoproteasome expression is lower, whereas a negative statistic indicates genes predicted to decrease when immunoproteasome expression is lower. For interpretability, the t statistic is equivalent to:

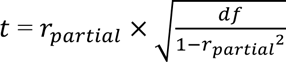

where *r_partial_* is the partial correlation between immunoproteasome expression and the target gene after covariate adjustment, and *df* is the residual degrees of freedom. The signed -*t* statistics were used for preranked GSEA and for plotting phagocytosis, lysosome, proteasome and mitochondrial gene trends.

#### Gene-set enrichment and module-level analyses

Pathway analyses were performed on ranked mouse and human differential-expression or regression results using GSEA from the R package WebGestaltR 0.4.6. Curated gene sets were obtained from Gene Ontology. Mouse gene lists were ranked by DESeq2 Wald statistics. Human immunoproteasome-low analyses were ranked by the signed -t statistic from the covariate-adjusted immunoproteasome-expression model.

Module scores were calculated from normalized expression values after within-dataset scaling and were analyzed with the same replicate-aware or donor-aware models used for gene-level analyses. Modules included constitutive proteasome, immunoproteasome catalytic subunits, PA28/proteasome activator genes, phagocytosis, phagosome maturation, lysosome, vesicle trafficking, actin remodeling, inflammatory signaling and nuclear-encoded mitochondrial genes. Module-level analyses were interpreted together with gene-level results rather than as independent confirmation of individual-gene effects.

#### Microglial state-marker and phagocytosis/cargo-processing analyses

Microglial state-marker and phagocytosis/cargo-processing genes were analyzed using the same mouse pseudobulk DESeq2 framework described above. Prespecified genes included homeostatic microglial markers, disease-associated microglial genes, phagocytic receptors and bridging molecules, cytoskeletal/cargo-engulfment genes, phagosome maturation genes, lysosomal genes and proteasome/immunoproteasome genes. Bar plots and heatmaps show DESeq2 log2 fold changes or signed Wald statistics from biological replicate-aware models.

For human snRNA-seq datasets, the same covariate-adjusted immunoproteasome-low regression framework was applied to microglial state-marker and phagocytosis/cargo-processing genes. The displayed value was the signed immunoproteasome-deficiency statistic (-*t*) from the model. The continuous immunoproteasome-expression model remained the primary analysis because it retained donor-level information and avoided dependence on an arbitrary threshold.

#### Multiple testing, software and reproducibility

Unless otherwise stated, *P* values from differential-expression, regression, pathway-enrichment, module-score and spatial analyses were adjusted using the Benjamini-Hochberg FDR method. The statistical unit for each model was defined according to the experimental design: mouse for mouse pseudobulk analyses, donor for human pseudobulk analyses and sample/donor for repeated spatial or predicted-proximity analyses. Transcriptomic analyses were performed in R 4.3.3 and, where applicable, Python 3.12.7. Software included Seurat, DoubletFinder, DESeq2, WebgestaltR or clusterProfiler, glmnet, lme4 or an equivalent mixed-model package, and visualization packages in R/Python. Generated mouse snRNA-seq data will be deposited in Synapse and reanalyzed human datasets will be cited with their AMP-AD/Synapse identifiers.

## Acknowledgements

We thank Tal Nuriel for helpful discussions and thoughtful insights throughout this study. We are grateful to Vilas Menon and Pallavi Gaur for their assistance with the ROSMAP dataset and related analyses. We thank Marc Diamond for generously providing the DS9 cells and the late Peter Davies for generously providing the PHF-1 antibody. We also thank Theresa Swayne from the Imaging Core for her guidance and technical support with live-cell imaging experiments. This study used the Confocal and Specialized Microscopy Shared Resource of the Herbert Irving Comprehensive Cancer Center at Columbia University, funded in part through NIH/NCI Cancer Center Support Grant P30CA013696. This work was supported by grants 1RF1NS143014 and R01AG070075 awarded to N.M. and R01AG075114 awarded to Q.W.

## Author contributions

N.M. conceptualized the study. N.M. wrote the original manuscript draft. M.S., S.J., S.M.W., S.S., M.Y., and Q.W. reviewed and edited the manuscript. N.M., Q.W. and M.Y. supervised the work. A.M.R. and M.S. contributed to Fig. 1 and Supplementary Fig. 1. D.E.L. and M.S. contributed to Fig. 2 and Supplementary Fig. 2. E.S., M.K, M.S., and M.Y. contributed to Fig. 3. A.M.R. contributed to Fig. 4. S.M.W., T.L. and Q.W. contributed to Fig. 5. S.J. contributed to Fig. 6 and Supplementary Figs. 3–10. S.J. and S.S. contributed to Fig. 7 and Supplementary Fig. 11.

